# Cerebral Organoids Uncover Mechanisms of Neural Activity Changes in Epileptogenesis

**DOI:** 10.1101/2025.08.26.672285

**Authors:** Sakurako Nagumo Wong, Michael Zabolocki, Oliver L. Eichmüller, Maryse A van ‘t Klooster, Marthe M. Priouret, Christian Krauditsch, Simone Krautberger, Julia Chu, Susana González-Granero, Lucas Barea Moya, Charles Fieseler, Segundo Jose Guzman, Daniel Reumann, Ramsey Najm, Jose Manuel García Verdugo, Mercedes F. Paredes, Manuel Zimmer, Maeike Zijlmans, Peter Jonas, Cedric Bardy, Nina Corsini, Jürgen A. Knoblich

## Abstract

Neurological disorders often originate from progressive brain network dysfunctions that start years before symptoms appear. How these changes emerge in the developing human brain remains elusive due to a lack of tractable model systems. Here, we show a cerebral organoid model of Tuberous Sclerosis Complex (TSC) that recapitulates hallmarks of epileptogenesis in vitro. We compare extracellular recordings of TSC organoids with intraoperative electrocorticography from TSC patients to reveal striking functional similarities, including high-frequency oscillations - an electrical biomarker for epileptogenic tissue. In TSC, a human-specific interneuron sub-type derived from the caudal ganglionic eminence drives network hyper-synchronization through increased spontaneous firing and altered excitability. Inhibiting overproliferation of its progenitors via long-term epidermal growth factor receptor inhibition prevented the onset of this pathological phenotype at functional and morphological levels. Our work shows that organoids allow mechanistic analysis of emerging neural network phenotypes, enabling anti-epileptogenic drug testing in a human brain development model.

## Introduction

Neurodevelopmental disorders originate in the prenatal brain and progress to debilitating conditions in early childhood. In epilepsy, seizures do not occur in isolation but are preceded by a progressive network dysfunction called epileptogenesis ^1,2^. Although animal models have provided key insights into epileptogenic network formation ^3^, human brain development contains unique neuronal subtypes ^4,5^, timing of circuit formation, and network cytoarchitecture ^6^, preventing direct translation of findings. Epileptogenesis can disrupt these developmental processes ^7^, making children particularly susceptible to epilepsy and seizure incidence highest in the first years of life ^8^. Anti-seizure medications primarily aim to suppress seizures after they occur without treating the cause and remain ineffective in approximately 37% of patients, despite decades of research ^9,10^. Understanding the process of human epileptogenesis and how distinct cell types contribute to pathological network formation is therefore critical for therapeutic innovation yet remains underexplored due to a lack of models that replicate human brain network development ^9,10^.

Cerebral organoids derived from human induced pluripotent stem-cells (hiPSCs) recapitulate key developmental milestones of the human brain in vitro, including human-specific cell types ^11–13^ and their specialized functions ^11^. Recent studies have shown that organoids can also generate functional networks capable of complex oscillatory network dynamics ^14–16^. Upon pharmacological perturbations, organoids exhibited synchronized oscillatory activity in high-frequency canonical bands ^16^, while others showed that low-frequency delta and theta-range oscillations can also emerge spontaneously, independent of stimulation ^14,15^. However, epileptogenic activity encompasses a diverse spectrum of signal properties and dynamic interactions between different frequency bands, reflecting underlying alterations in cellular mechanisms ^17^. Whether organoids can spontaneously generate epileptogenic activity analogous to that observed in children with epilepsy remains underexplored. Further, the extent to which cerebral organoids can model epileptogenesis remains unclear.

We have recently recapitulated the histological abnormalities of the genetic epilepsy syndrome Tuberous Sclerosis Complex (TSC) in a patient-derived organoid model ^18^. TSC is caused by mutations in *TSC1* or *TSC2*, resulting in drug-resistant epilepsy (DRE) ^19^, with seizures occurring mostly during infancy in up to 90% of patients ^20^. The morphological hallmarks of TSC include benign brain tumors and cortical tubers that emerge during brain development ^21^. Cortical tubers, containing dysmorphic neurons (DNs) ^22^, have been identified as the foci of epileptic activity ^23–25^. Importantly, surgical resection of these tubers has been shown to reduce both seizure frequency and severity ^26^. Using TSC organoids, we uncovered that the morphological alterations are initiated by an over-proliferation of a human-specific interneuron progenitor population that originate from the caudal ganglionic eminence (CGE) at mid-gestation, referred to as caudal late interneuron progenitor (CLIP) cells ^18^. CLIP cells gave rise to dysmorphic GABAergic interneurons ^18^, but whether they participate in the network and lead to epileptogenic activity remains unclear. In TSC patients, intra-operative electrocorticography (ioECoG) recordings in tuber regions during interictal periods revealed an increase in interictal epileptiform discharges (IEDs) as a marker for pathological networks ^24,27^. Moreover, high frequency oscillations (HFOs) within ripple (80 – 250 Hz) and fast ripple (250 – 500 Hz) frequency bands have recently emerged as a highly predictive interictal biomarker for the detection and resection of epileptogenic tissue ^28,29^. While surgical resection is not always feasible ^30^, analysis of these electrophysiological biomarkers at a higher temporal resolution provides insights into the underlying dysregulated network and the genesis of seizure activity ^27,31^.

In this study we combined calcium imaging and extracellular recordings to systematically investigate spontaneous epileptogenic activity in long-term cultures of patient-derived TSC organoids. Compared to isogenic controls, TSC organoids exhibited hypersynchronous firing, dysregulated oscillatory activity, and excitotoxic morphological changes. Strikingly, HFOs emerged in TSC organoids, closely resembling those measured in ioECoG recordings from TSC patients. Single-unit spike analysis revealed that HFO events in control organoids were accompanied by staggered, oscillatory firing between excitatory and interneuron populations, suggestive of excitatory-inhibitory feedback ^32^. This coordinated activity was lost in TSC organoids due to shifts in firing that were associated with the emergence of network hyper-synchronization. Long-term inhibition of the epidermal growth factor receptor (EGFR) reduced both the pathological electrophysiological and morphological phenotypes, including dysmorphic interneurons. Together, our study demonstrates the utility of cerebral organoids as a human-specific developmental model for identifying the responsible mechanisms underlying pathological network formation during epileptogenesis and applying anti-epileptogenic drug testing.

## Results

### TSC organoids recapitulate aberrant hyperexcitable network dynamics

Cerebral organoids generated from TSC patients can recapitulate the formation of tuber-like structures ^18^, a key diagnostic hallmark of the disease. In TSC organoids, tuber development is initiated by interneurons that are generated by CLIP cells starting after around four months of culture ^18^. To investigate whether the organoid model can also recreate disease-specific neural network activity, we used functional calcium imaging and extracellular recordings (Methods, Figure 1A). Organoids derived from two TSC patients and their isogenic controls were cultured in a low-nutrient medium for an average of six months ^33^. Organoid integrity and cell type composition were verified by immunostaining for common lineage markers to ensure the presence of both excitatory and inhibitory neuronal populations (Figures S1A and S1B). To identify differences in network activity between control and TSC organoids, we inserted a construct expressing GCaMP6s under the human synapsin promoter into the AAVS1 locus (Methods, Figure 1A). Synchronous and recurring population calcium network events were quantified at approximately six months (Figure 1B) across a total of 1007 and 1095 cells from 53 and 47 organoids, respectively. Comparing calcium transients in individual cells, TSC organoids displayed significantly more frequent network events, indicating elevated network activity (Figures 1B to 1C, Video S1). Next, we performed acute extracellular recordings with 64-channel Cambridge Neurotech silicon probes to measure spontaneous neuronal activity with higher temporal resolution (Figure 1A; Methods). Coordinated action potential firing was recorded across multiple electrodes (Figure 1D top), referred to as multi-unit activity (MUA). During brief periods in extracellular recordings, MUA spontaneously synchronized across neighboring channels, herein referred to as ‘network events’ (Figure 1D). To evaluate the role of both synaptic and gap-junction-mediated transmission on coordinated network event generation, we applied different pharmacological interventions. The application of 200 µM carbenoxolone (CBX), inhibiting gap-junctions, did not affect network event activity or MUA firing rates (Figures 1E, S1C and S1G). In contrast, 0.5 µM tetrodotoxin (TTX) completely abolished network event activity and MUA firing (Figures 1E, S1E, and S1G) ^15,16^. Notably, 10 µM bath applications of the AMPA-receptor-antagonist CNQX had similar effects (Figures 1E, S1D, and S1G), validating the role of glutamatergic synaptic transmission in synchronized network activity. Considering that during neurodevelopment, post-synaptic responses of GABAergic transmission switch from depolarizing to hyperpolarizing inhibitory effects due to changes in chloride homeostasis ^34^, we next evaluated the impact of GABA neurotransmission on network event activity. Indeed, applications of 20 µM GABA during extracellular recordings abolished all synchronized network activity and MUA firing, which partially recovered upon washout (Figures 1F, S1F and S1H). Overall, these results indicate that synchronized network events in cerebral organoids rely on synaptic activity and require both excitatory glutamatergic and inhibitory GABAergic transmission.

**Fig. 1.**
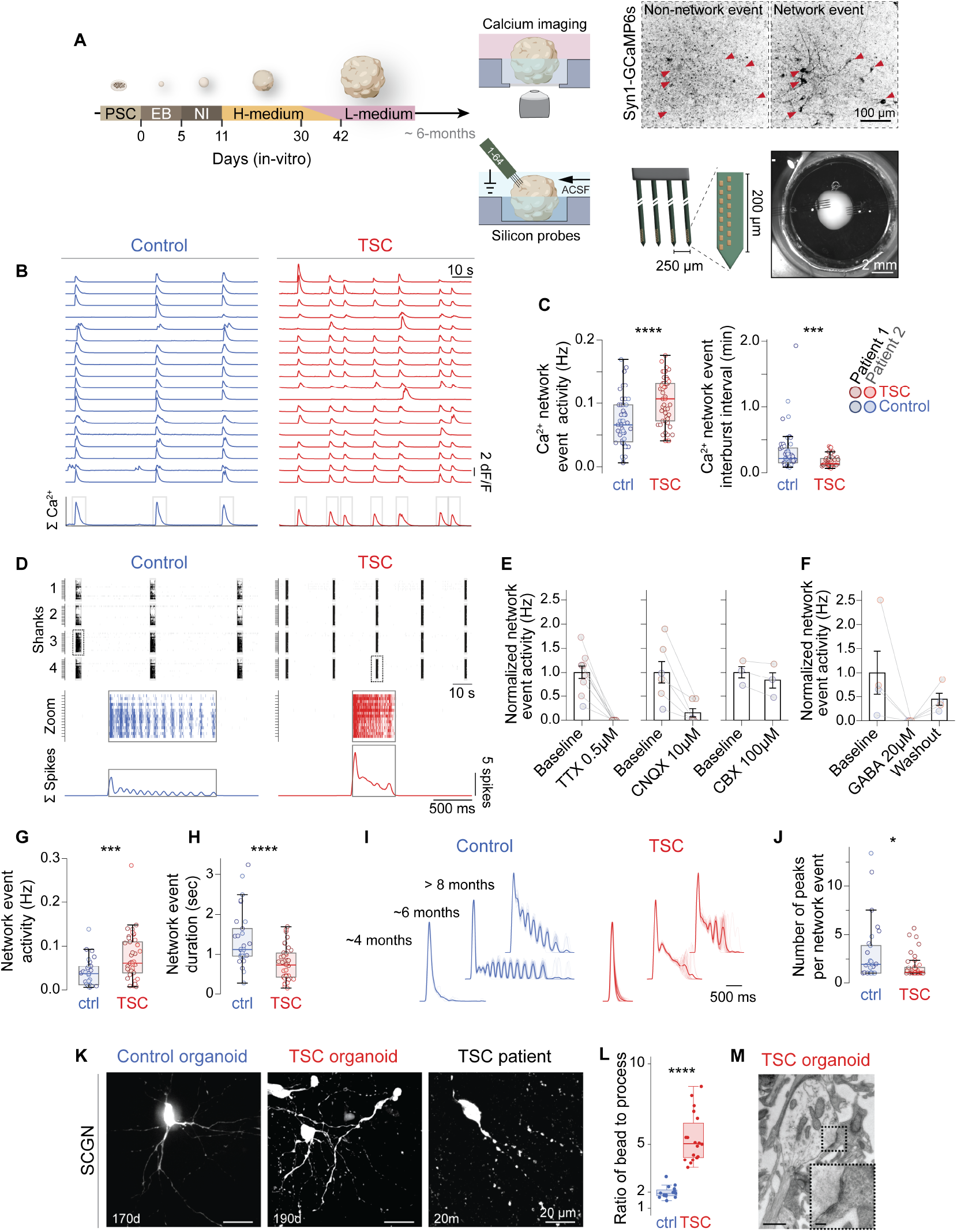
Hyperexcitability impairs network function in TSC. (A) Schematic describing the generation of cerebral organoids from control and TSC hiPSC cells, first in high-nutrient media then matured in low-nutrient media (left), along with the recording of spontaneous extracellular signals (bottom right) and calcium activity in organoids (top right). Representative snapshots during GCaMP imaging before and after calcium network events are highlighted (top right). Neuron cell-bodies are highlighted (red arrows). (B) Representative traces of Ca^2+^ transients from 18 cells in control and TSC organoids at approximately 6-months, along with their summed activity (bottom). Detected calcium network events are highlighted in grey boxes. (C) Quantification of calcium network event activity frequency and inter-burst interval (n = 53 for control and n = 47 for TSC organoids from 7 differentiation experiments of four hiPSC cell-lines). (D) From top to bottom: representative raster plots of multi-unit spike activity from control and TSC organoids, followed by a zoom in of multi-unit spike activity from an isolated shank (box), and the sum of neuronal spikes (bottom). (E and F) Pharmacological perturbations of network events following 0.5 µM TTX, 10 µM CNQX and 200 µM CBX applications (E), and 20 µM GABA (F) in control and TSC organoids. Application of GABA was partially reversed during washout in organoids. Data are shown as mean ± SEM. (G and H) Quantification of network event activity (G) and duration (H) from control and TSC organoids (n = 29 for control from 11 differentiation experiments, and n = 41 for TSC organoids from 13 differentiation experiments of four hiPSC cell lines). (I) Representative network events from single control and TSC organoids at approximately 4, 6, and above 8 months in vitro revealing increasing nested oscillatory subpeaks in network maturation. Isolated network events are overlaid and normalized to maximum peak amplitudes. (J) Quantification of the number of peaks per network event revealing reduced nested oscillatory subpeaks in TSC organoids (n = 29 for control from 11 differentiation experiments, and n = 41 for TSC organoids from 13 differentiation experiments of four hiPSC cell lines). (K) Immunofluorescence images of CGE interneurons marked with SCGN in control-, TSC organoids, and TSC patient tissue identifies excitotoxicity with dendritic beading in TSC organoids and patients. (L) Increased dendritic beading to process length ratios are identified in TSC. P-values calculated using Wilcoxon signed-rank test (control, n = 15; TSC, n = 20 processes from 4 organoids; P < 0.0001). (M) Electron microscopy (EM) of TSC organoids identifies damaged and enlarged postsynaptic beading on dendrites. Scale bars: overview 1 µm, zoom 200 nm. Boxplots shown in (C), (G), (H), (J): center, median; lower hinge, 25% quantile; upper hinge, 75% quantile; whiskers ± 1.5 x interquartile range. Data shown in (E) and (F) are presented as mean ± SEM. *P<0.05; ***P<0.001; ****P<0.0001.

Detection of synchronous network events during extracellular recordings allowed us to quantify differences between control and TSC patient organoids. In TSC, network event durations decreased from 1.4 s (+/-0.13 s, n = 29 organoids) to 0.76 s (+/-0.067 s, n = 41 organoids, P < 0.0001), while the frequency of events increased from 0.039 Hz (± 0.0059 Hz) to 0.076 Hz (± 0.0084 Hz, P = 0.0009, Figures 1G and 1H). Of note, changes in network event durations and event frequencies suggest differences in the underlying ratios of excitatory and inhibitory synaptic charge ^35^. To further evaluate this, we tested whether organoids could generate nested oscillatory dynamics during network events, which are driven by balanced synaptic excitatory and inhibitory inputs as the network develops ^36,37^. Indeed, investigating synchronized spiking activity in control organoids revealed reverberant oscillatory peaks during network events in control organoids (Figures 1D and S1I). A nested periodic oscillatory pattern occurred 200-500 milliseconds after initial network activation, coinciding with low-frequency oscillations (∼4-8 Hz) in single-channel local field potentials (LFPs) (Figures 1D and S1I). Moreover, power above ∼15 Hz rapidly diminished following the peak response in control organoids, whereas in TSC organoids, power remained sustained (Figure S1I). Consistent with previous studies evaluating nested network-event oscillatory dynamics using multi-electrode array recordings ^15^, its periodicity became less frequent after 6-months (Figures 1I). TSC organoids, in contrast, failed to replicate oscillatory patterns during network events, even after ∼6-months (P = 0.036; Figures 1I, 1J, and S1I). In addition to impaired maturation, hyperexcitable network dynamics in epileptogenic regions of patients are often accompanied by abnormal cellular morphologies reflecting excitotoxicity ^38,39^. To test whether this is recapitulated in TSC organoids, we examined the dendritic morphologies of SCGN positive, CLIP cell-derived interneurons that are altered in TSC ^18^. Dendrites identified by MAP2 staining and transmission electron microscopy displayed focal swellings (Figures 1K to 1M, and S1J and S1K) that were previously associated with excitotoxicity ^40,41^. Thus, TSC organoids exhibit impaired maturation of neural network activity and signs of excitotoxicity, which may underlie the elevated occurrence of synchronous network events and give rise to a pathological network.

### Pathological HFOs mark epileptogenic tissue in children with TSC

The increase in MUA activity and altered network dynamics might indicate that TSC organoids can recapitulate the changes in neural networks during epileptogenesis. To establish a reference for distinguishing between physiological and pathological network events, we analyzed data from two children with TSC ^29^. Specifically, we investigated pathological network activity through intraoperative electrocorticography (ioECoG) recordings before and after the surgical removal of epileptogenic cortical tubers (Figures 2A, 2B, and S2). HFOs both within the ripple (80 – 250 Hz) and fast ripple (250 – 500 Hz) range have previously been employed as biomarkers to distinguish physiological and pathological network activity ^27,42–44^. However, differences in time-frequency signal features between pathological and physiological HFOs remain inconsistent, largely due to the absence of a stan-dardized definition and detection methodology across disciplines ^17^. To address this, putative HFOs were detected using automated HFO detectors and then visually inspected by an experienced clinical examiner (Figure 2C, Methods). All accepted HFO events adhered to clear clinical-based criteria (Methods) to ensure robust comparisons between cortical tubers and physiological and functional cortical regions. Indeed, measurements in the cortical areas overlying tuber lesions revealed pathological events that had characteristically coinciding ripples, fast ripples, and interictal epileptiform discharges (IEDs, 20 - 80 Hz; Figure 2C). In contrast, recordings from unaffected central motor areas, which are indispensable for normal brain function ^45^, identified events classified as physiological HFOs (Figure 2C). When comparing physiological and pathological ripple activity, epilepti-form events were characterized by significantly shorter duration, higher amplitude, and higher intrinsic disorganization, measured as ripple entropy (Figure 2D). Notably, ioECoG-tailored surgery based on pathological HFOs and identifying the margins of the tuber tissue resulted in long-term seizure freedom in both children (Figures 2A, S2 and Table S1). This indicates that the intrinsic properties of pathological HFOs reliably identified epileptogenic regions (Figures 2A and S2), providing a crucial reference for interpreting and validating the hyperexcitability phenotypes observed in the TSC organoids.

**Fig. 2.**
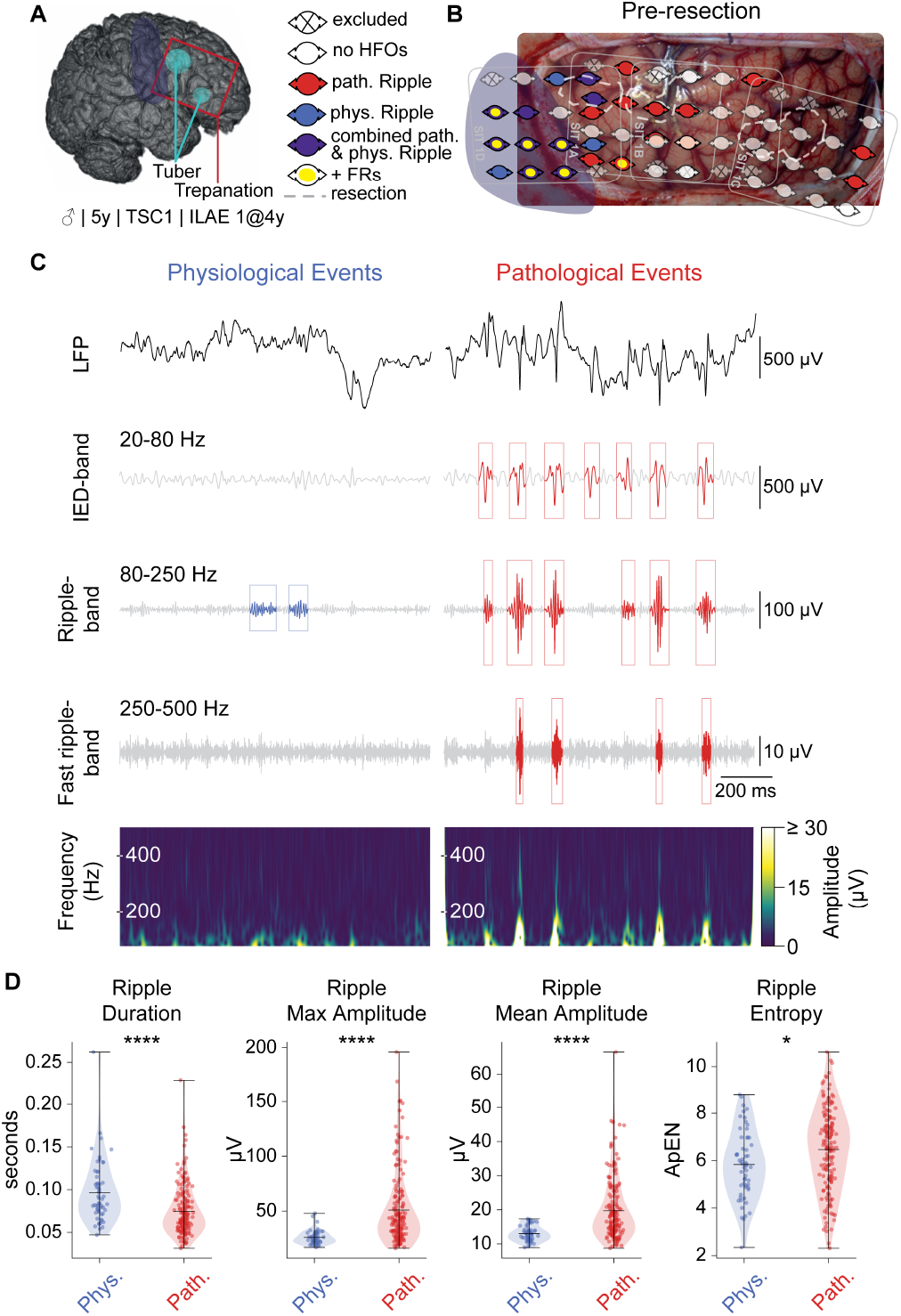
Pathological HFOs mark epileptogenic tissue in TSC patients. (A) 3D MRI rendering depicting two cortical tubers in the right frontal lobe (light blue) of a TSC patient (Pat #1). A schematic of the trepanation window is shown in red. The eloquent central motor region is marked in dark blue. (B) ioECoG grid electrode placement in multiple positions covering the frontal tubers. Physiological events were recorded in the central motor frontal areas, occasionally on electrodes overlapping with pathological ripples and fast ripples. (C) Representative physiological (blue) and pathological (red) HFOs. LFP (top), IEDs (20–80 Hz), ripple (80–250 Hz), and fast-ripple signal bands (250–500 Hz) with a corresponding time–frequency spectrogram (bottom). (D) Ripple-feature quantifications across four pre-resection grid positions in this TSC patient. Physiological (n = 57) vs pathological (n = 165) ripples (Mann–Whitney: duration P < 0.0001; max amplitude P < 0.0001; mean amplitude P < 0.0001; entropy P = 0.0107). ApEN, approximate entropy. Boxplots: center = median; lower hinge = 25% quantile; upper hinge = 75% quantile; whiskers = ±1.5 interquartile range.

### Epileptogenic HFOs are prevalent in TSC organoids

To evaluate whether TSC organoids can recapitulate the pathological network activity found in TSC patients, we first isolated the IED, ripple, and fast-ripple frequency bands from high amplitude channels in control and TSC organoids (Methods, Figures S3A and S3B). HFO and IED events were separately detected in both the time and frequency-domains using automated custom computational pipelines that adhere to standard clinical detection practices for patient ioECoG recordings (Methods, Figures S3A to S3D). Visual inspections of all detected events were performed in-line with published classification protocols to manually exclude false-positives (Methods, Figures S3C and S3D). Acute recordings in both control and TSC organoids revealed spontaneously generated HFOs with a spectrum of morphological features (Figures 3A and 3B).

**Fig. 3.**
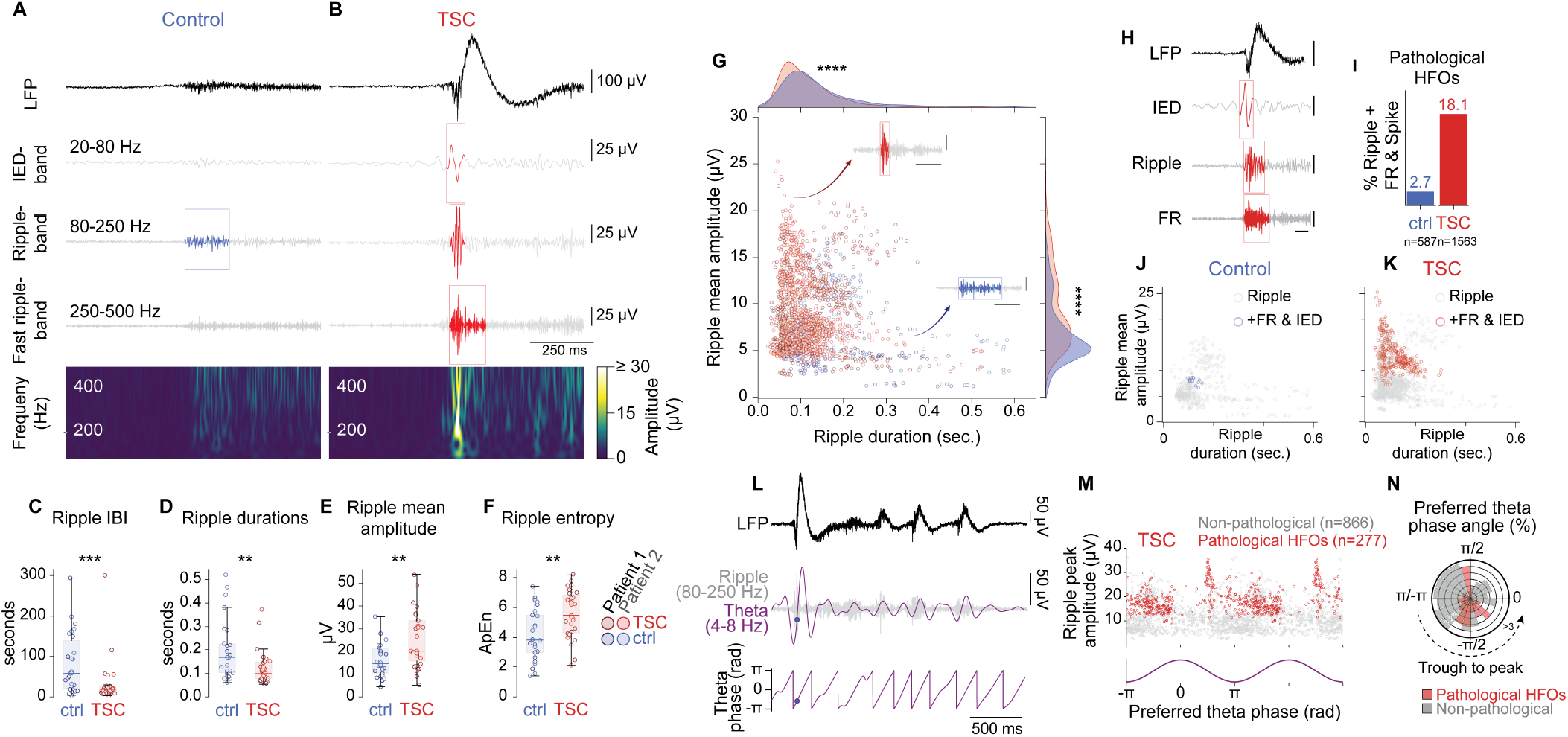
Epileptogenic HFOs emerge in TSC organoids. (A) and (B), Top: Representative LFP and band filtered signals from control and TSC organoids. Timestamps at detected IED, ripple, and fast ripple events are marked with boxes (control, blue; TSC, red). From top to bottom: LFP, IED, ripple, fast ripple, time–frequency spectrogram. Color bar reflects the signal amplitude (µV). (C), (D), (E), and (F) Quantified ripple signal properties: inter-burst interval (C, IBI, P = 0.0009), duration (D, P = 0.0073), amplitude (E, P = 0.0019), and entropy (F, P = 0.0036) across baseline periods (control, n = 26 organoids from 11 differentiation experiments; TSC, n = 31 organoids from 11 differentiation experiments; ApEN, approximate entropy). Boxplots shown as: center, median; lower hinge, 25% quantile; upper hinge, 75% quantile; whiskers *±* 1.5 *×* interquartile range. (G) Joint plot illustrating the distributions of the mean amplitudes and durations of single ripples in control (n = 587) and TSC organoids (n = 1563 ripples). Insets highlight representative ripple waveforms corresponding to selected data points (arrows). Kolmogorov–Smirnov tests were performed between pooled datasets, revealing significant differences in ripple mean duration (P *<* 0.0001) and mean amplitude (P *<* 0.0001). Inset scale bars: 25 µV (ripple, vertical) and 250 ms (time, horizontal). (H) Representative IED and fast ripple coinciding at an isolated ripple event. From top to bottom: LFP, IED, ripple-, fast ripple-band filtered trace. (I), (J), and (K), Comparison of coinciding ripples with IEDs and fast ripples (FRs) depicting proportion of all events (I), and the distribution across the joint plot of mean ripple amplitude and duration in single ripples from control (J, total n = 587, +FR & IED n = 16) and TSC organoids (K, n = 1563, +FR & IED n = 283). The colors in the joint plot identify ripple events coinciding with both IEDs and fast ripples. (L) Representative example of phase–amplitude coupling analysis from an LFP recording (top). Theta oscillations (purple, middle) and ripple-band activity (gray, middle) are isolated around ripple events. The preferred theta phase of ripple peaks (circles) is determined to assess phase– amplitude coupling angle preference (bottom). (M) Distribution of ripple peak amplitudes across the theta phase cycle (bottom, purple). Pathological high-frequency oscillations (HFOs) in TSC (red) are identified based on high amplitude ripples and their co-occurrence with IEDs and fast ripples. Pathological HFOs preferentially occur during the trough-to-peak phase of theta (*−π* to 0). Non-pathological ripples (gray) show a more uniform distribution with no coupling preference. For visualization, one theta phase cycle is repeated. (N) Rose plot showing the preferred theta phase angle distribution of ripples from (M). Pathological HFOs in TSC (red) predominantly couple to the trough-to-peak phase of theta, whereas non-pathological ripples (gray) exhibit weaker phase preference. For (C) to (F), colors correspond to patient-derived iPSC cell lines.

Ripples occurred in shorter intervals in TSC organoids compared to controls, supporting a hyperexcitability phenotype (Figure 3C), and were associated with larger field potentials deflections that propagated across channels (Figure S3E). Remarkably, TSC organoid ripples also displayed increased amplitudes, shorter durations, and higher entropy (Figures 3D to 3F and S3F), like pathological activities at local epileptogenic cortical tubers in children with TSC (Figure 2D) ^29^. Subsequently, HFOs frequently appeared with irregular amplitudes and morphologies in TSC organoids (Figure 3B), also mirroring the characteristic HFOs observed in epileptogenic tissue ^46^. Thus, our data suggest that networks in TSC could replicate pathological network activities of cortical tubers.

HFOs vary in amplitude and duration along a spectrum, reflecting the degree of underlying spike activity (Figures 2D, 3G, and ref. ^47^). However, ripple occurrence alone is not predictive of epileptogenic tissue (Figure 2) ^48^. Therefore, to guide epileptogenic tissue resections in children with TSC (Figure 2C) ^49^, ripples that co-localize with fast ripples, IEDs, and ‘spike’ like morphologies in field potentials (hereafter referred to as ‘pathological HFOs’) are commonly used instead ^50^. To evaluate whether this complex set of features is recapitulated in the organoid model, we tested to what extent ripples occurred together with IEDs and fast ripples (Figures 3H to 3K). While ripples and fast ripples were detected as before (Figures 3A, S3C, S3D), IEDs were identified based on a published method modified for organoids (Methods, Figure S3D and ref. ^51^). In control organoids, only a small frac-tion of HFOs coincided with IED-like events and fast ripples (2.7%), while in TSC organoids this was drastically increased (18.1%; Figures 3G to 3I) and mainly included characteristic high-amplitude pathological ripples (Figures 3J and 3K). Moreover, fast ripples detected in TSC organoids exhibited hallmark pathological features, occurring more frequently with higher amplitudes and shorter durations (Figures S3G and S3H). Taken together, TSC organoids replicate abnormalities of interictal pathological HFOs, including their interactions with other epileptogenic biomarkers. Thus, the intrinsic properties shared between the organoid model and pathological HFOs from patients suggest that TSC organoids develop similar pathological network properties.

In epilepsy patients, local alterations in epileptogenic regions lead to changes in the modulation of high-frequency amplitudes by low-frequency phases, a phenomenon referred to as cross-frequency phase-amplitude coupling (PAC) ^52–55^. Specifically, HFOs are observed more often during distinct phases of low-frequency oscillations. Quantifications of PAC between low-frequency theta (4-8 Hz) and high-frequency ripple power (80-250 Hz) revealed stronger coupling between ripples and theta-oscillations in TSC organoids when compared to control organoids (Figures 3L and 3M, S3I and S3J). This finding mirrors network changes in epilepsy patients, where theta-ripple coupling was strongest during seizure evolution ^52^. In patients, HFOs classified as pathological couple to distinct phases of theta activity and occur more frequently during trough-peak transitions ^54,55^. We therefore evaluated the phase-preference of theta-ripple coupling in organoids during HFO events that were classified non-pathological or pathological. Strikingly, the peak amplitudes of pathological HFOs observed in TSC organoids concentrated at trough-peak phases of theta, near -π/2 (Figures 3M and 3N). In contrast, those HFO events in TSC organoids that are classified as non-pathological displayed a diverse theta phase-preference in TSC, reflecting low theta-ripple coupling (Figures 3M and 3N). Thus, larger ripple amplitudes detected in pathological HFOs more consistently coupled to theta oscillations at trough-peak phases. Taken together, TSC organoids recapitulated major hallmarks of epileptogenic networks as pathological events containing high-amplitude HFOs and IEDs, as well as phase-preferences to pathological theta phases.

### Dysregulated firing of neuronal subtypes shapes pathological activity in TSC networks

Compared to ioECoG recordings in patients, extracellular recordings in organoids using high-density silicon probes offer superior temporal resolution and spatial density, enabling more precise characterization of neural activity. This allowed us to evaluate the altered firing patterns underlying the changes in HFOs seen in TSC organoids. To this end, we used a unified spike-sorting pipeline ^56^ to infer the biophysical characteristics of single neurons ^57^. Spike-units were extracted from control and TSC organoids, yielding a total of 207 neurons (Methods, Figures 4A and S4A). We isolated the maximum amplitude waveforms and applied an unsuper-vised clustering method ^58,59^to group units based on extracellular waveform features (Methods, Figures 4A, S4B and C). This analysis yielded four distinct groups with differing biophysical and firing properties (Methods, Figures 4A to 4D, and S4C to S4G). Waveforms and autocorrelograms (ACGs) corresponding to these clusters reflected unique putative cell types (Figure 4C) that a random forest classifier could predict with a precision of up to 100% (Methods, Figures S4D). This suggests that the clusters of extracellular waveforms detected in cerebral organoids correspond to functionally distinct neuron classes, allowing us to draw conclusions on the network composition.

**Fig. 4.**
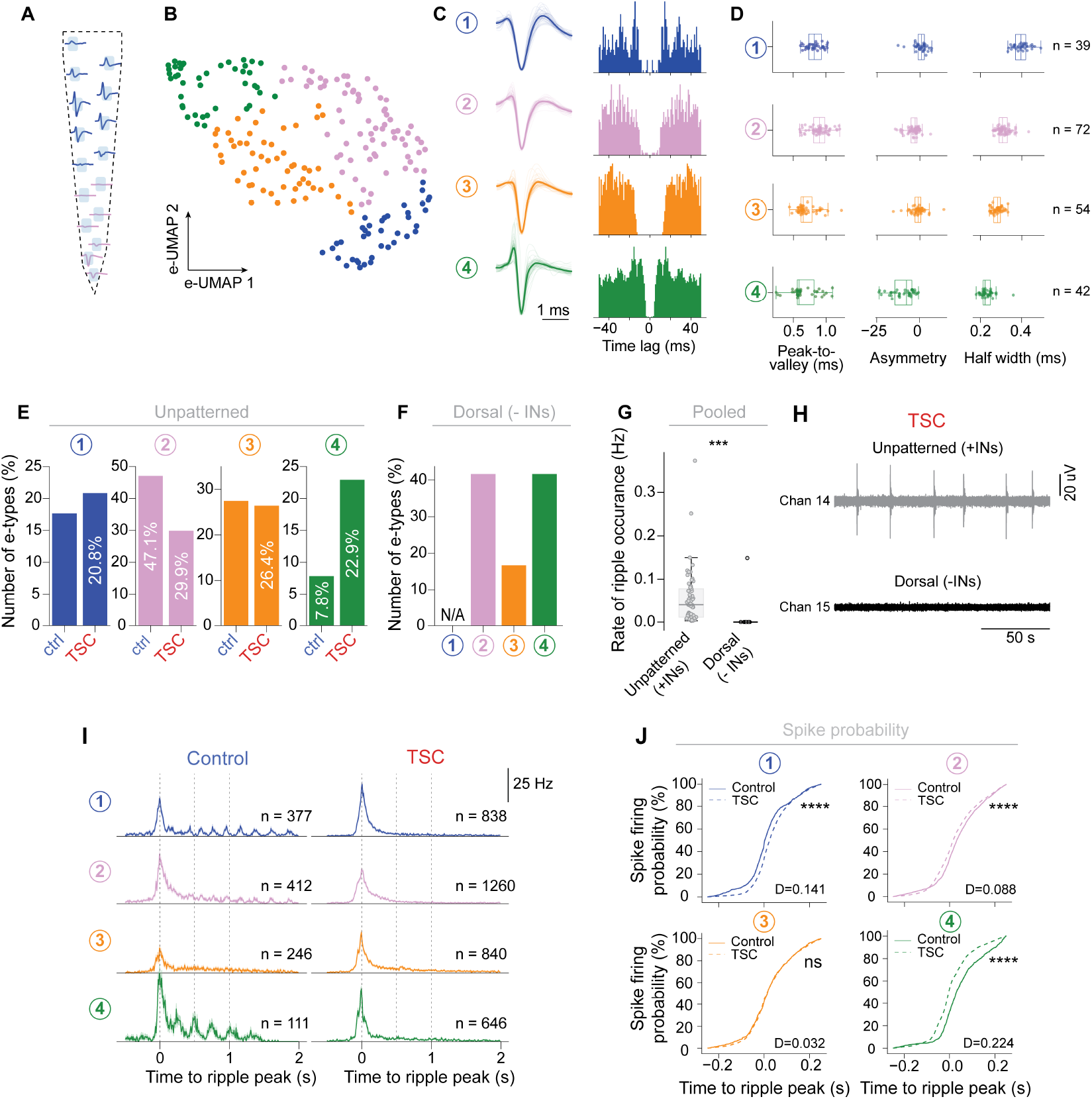
CGE interneurons are dysregulated in TSC networks. (A) Example single 16-channel Cambridge Neurotech silicon probe shank resolving neuronal spiking in a TSC organoid. (B) Uniform manifold approximation and projection (UMAP) dimensionality reduction based on normalized spike waveforms, extracted from 207 neurons across control (n = 17 organoids from 8 differentiation experiments) and TSC organoids (n = 35 organoids from 14 differentiation experiments). Data point colors mark waveform clusters (see Methods). (C) Average waveforms of each cluster (left) overlaid on traces of all individual waveforms, illustrating cluster-specific spike shapes. Representative autocorrelograms (right, color scheme as in b) display the temporal firing patterns for each cluster. (D) Peak-to-valley duration, waveform asymmetry and half-width values for each spike unit witin each waveform cluster. Corresponding n-values are displayed. (E) and (F) Spike waveform cluster contributions to control and TSC organoids in unpatterned (control: n = 51 cells; TSC: n = 144 cells) and dorsal protocols (TSC: n = 12 cells from 4 organoids). (G) Quantification of ripple events occurance rates during baseline periods in dorsal-patterned cerebral organoids lacking CGE-derived interneurons and pooled unpatterned organoids containing interneurons. P-values calculated using Wilcoxon signed-rank test (unpatterned, n = 57 organoids across 22 differentiation experiments; dorsal, n = 8 organoids across 3 differentiation experiments; P = 0.0005). Data represented as mean +/-SEM. (H) Representative single-channel ripple signal traces from unpatterned and dorsal TSC organoids. (I) Perievent spike-time distributions and (J) spike firing probability for each spike waveform across ripple events in control (n = 17) and TSC organoids (n = 31) organoids. Mean firing rates of units are shown for cluster 1 (control, n = 377 events from 7 cells; TSC, n = 838 events from 20 cells), waveform cluster 2 (control, n = 412 events from 20 cells; TSC, n = 1260 events from 34 cells), waveform cluster 3 (control, n = 246 events from 12 cells; TSC, n = 840 events from 26 cells), and waveform cluster 4 (control, n = 111 events from 4 cells; TSC, n =646 events from 15 cells). Color scheme matches (B). Data represented as mean +/-SEM. Statistics in (J) calculated using the Kolmogorov-Smirnov test. *P<0.05; ***P<0.001; ****P<0.0001. The Kolmogorov-Smirnov statistics (‘D’) are shown for each spike waveform type between conditions.

The identification of distinct spike types enabled us to compare spike type proportions between control and TSC organoids. In TSC, waveform cluster 2 was reduced, whereas clusters 1 and 4 were enriched (Figures 4E, S4F and S4G). Cluster 1 exhibited an increased spontaneous burst index while cluster 4 is characterized by an increase in firing rate and reduced half-width and peak-to-valley durations (Figure 4D and S4C and S4E). Both features highlight a potential role of these spike units and the corresponding types of neurons in pathological network synchrony ^17^, as increased firing rate and burst-index spikes features have been observed during interictal periods in human epilepsy ^60^. To identify which cell types are responsible for the individual waveforms, we generated dorsalized organoids in which the number of interneurons is drastically reduced ^18^. Dorsalization of organoids and a decrease of interneurons was verified by staining for the CGE interneuron marker secretagogin (SCGN; Figure S4H). Spike-sorting and waveform clustering of these dorsalized organoids revealed the persistence of clusters 2, 3, and 4 while cluster 1 was absent, suggesting that this waveform class corresponded to an interneuron cluster. Moreover, cluster 1 exhibited the highest firing rates and B-peak amplitudes – features characteristic of interneuron extracellular waveforms (Figure S4E) ^61,62^. Notably, half-width values were broader than previously characterized narrow-spiking parvalbumin (PV) interneurons ^63^, suggesting this waveform cluster represented a wide-spiking CGE interneuron subtype, likely originating from CLIP cells previously found in cortical tubers ^18^. In contrast, clusters 2, 3, and 4 persisted in dorsalized organoids, representing putative excitatory neurons, where clusters 2 and 3 may retain immature electrophysiological properties ^64^ (Figure S4E). Compared to unpat-terned organoids, these dorsalized organoids exhibited a significant reduction in spontaneous HFOs (Figures 4G and 4H, and S4H) and 87.5% of them displayed no detectable ripple events (Figure S4J), underscoring the requirement of CLIP cell-derived interneurons in generating pathological HFOs. Thus, clustering of extracellular waveforms allowed us to draw conclusions on the putative excitatory and interneuron populations that may be responsible for the changes in neural network activity observed in TSC.

By identifying unique waveform clusters, we asked how individual spike types align with peak HFO amplitudes to understand how different contributions of neuronal subtypes lead to network synchronization and epileptogenic activity. To test this, we analyzed the firing rates for each waveform relative to the peak amplitude of detected HFO events (Methods, Figure 4I). In control organoids, all waveforms showed a longer firing period, an observation consistent with the increased HFO duration seen compared to TSC organoids (Figure 4I and Figures 3A, 3B, and 3D). Interestingly, we observed a pattern of oscillatory firing between putative interneuron (cluster 1) and excitatory neurons (clusters 2 and 4) across several control organoids, that might be indicative of nested oscillatory dynamics observed during network events (Figures 4I, 1I, and S1I). After the ripple peak, putative excitatory neurons fired first, followed by staggered firing from putative interneurons that shaped the elongated ripple. In contrast, no oscillatory subpeak formation was observed in TSC organoids, as all waveforms fired synchronously around the ripple peak (t=0). Effective putative interneuron (cluster 1) firing was 27.32 ± 2.44 Hz (n = 377 events from 7 cells) in control and 37.71 ± 2.16 Hz (n = 838 events from 20 cells) in TSC organoids, shifts in firing probability, from 53.20% of spikes firing after the ripple peak in marking heightened firing frequencies during HFO events (Figure 4I). This was accompanied by control to 66.20% in TSC organoids (Figure 4J). This increase and shift in interneurons firing after the peak might prevent firing from other clusters and explain the shorter total firing period, we observe in TSC organoids. Spikes assigned as putative excitatory neurons (clusters 2 and 4), however, fired before the ripple peak in 41.25% (cluster 2) or 53.23% (cluster 4) of cases in TSC organoids. This was accompanied by decreased firing frequencies in TSC relative to control organoids in both cluster 2 (control: 35.68 ± 2.84 Hz, n = 412 events from 20 cells; TSC: 24.68 ± 1.30 Hz, n = 1260 events from 34 cells) and cluster 4 (control: 49.55 ± 5.96 Hz, n = 111 events from 4 cells; TSC: 38.24 ± 2.29 Hz, n = 646 events from 15 cells). None of the neurons in cluster 3 displayed a preferential firing time (Figure 4J), suggesting minimal contribution, although the maximum firing frequencies varied (control: 17.89 ± 2.70, n = 246 events from 12 cells; TSC: 29.17 ± 1.94 Hz, 40 events from 26 cells). Together, these results suggest asymmetric firing of neuronal subtypes during HFOs, with enhanced network synchronization in TSC organoids driven by increased interneuron firing and altered excitatory-interneuron timing. These changes may lead to the formation of short duration, high amplitude pathological events that are also present in the epileptogenic regions of TSC patients.

### TSC organoids can uncover anti-epileptogenic drug effects

The changes in firing patterns of excitatory and inhibitory populations in TSC organoids led us to speculate whether the dysmorphic GABAergic interneurons previously found in tuber-like regions could result in hyperactive network dynamics, ultimately causing downstream pathological network activity ^18^. To this end, TSC organoids previously subject to extracellular recordings were analyzed, confirming the presence of MAP2-positive neurons with high phosphorylated-S6 (PS6) protein levels, a marker for mTOR activation, and pathological cell types (Figure 5A and ref. ^21^). Interestingly, dysmorphic neurons had MAP2+ processes, further supporting the inclusion of pathological cell types in the network, as evidenced by synapses observed in electron microscopy (Figure 1M). As the epidermal growth factor receptor (EGFR)-inhibitor Afatinib has proven effective in reducing the proliferation of CLIP cells that give rise to dysmorphic interneurons ^18^, we tested whether targeting the EGFR pathway could thereby inhibit epileptogenesis. Afatinib was added long term to organoid cultures from three months (Figure 5B) when CLIP cells first start to proliferate in TSC ^18^ and before any electrical phenotypes have developed. Recordings with silicon probes at approximately 7 months captured a developmental stage where major epileptogenic changes would have already occurred. Comparison of pathological events in untreated TSC organoids with long term Afatinib-treated TSC organoids revealed a decrease in the number of fast ripples (Figures 5C and 5D), as well as a significant increase in the mean duration of each event (Figure 5E), revealing characteristics closer to non-pathological HFO events in control organoids (Figure 3). To confirm that these changes in fast ripple characteristics were due to the reduction of dysmorphic interneurons, we quantified the number of GAD67+ neurons with dysmorphic morphology in untreated and Afatinib-treated TSC organoids (Figures 5F and 5G). The long-term treatment of Afatinib led to a striking reduction in the number of dysmorphic interneurons, suggesting that long term treatments affecting epileptogenesis at the network-level is also reflected in changes in morphological phenotypes. Subsequently, as a downstream effect of long-term Afatinib treatment in TSC organoids, we observed that interneurons exhibited healthy morphology without focal dendritic swellings, suggesting a reduction in excitotoxicity and reversal in hyper-excitable network dynamics (Figure 5H). Thus, these results suggest that dysmorphic interneurons contribute to the generation of epileptogenic activity in TSC, and inhibiting their proliferation in the developing brain may offer a promising strategy for anti-epileptogenic treatments.

**Fig. 5.**
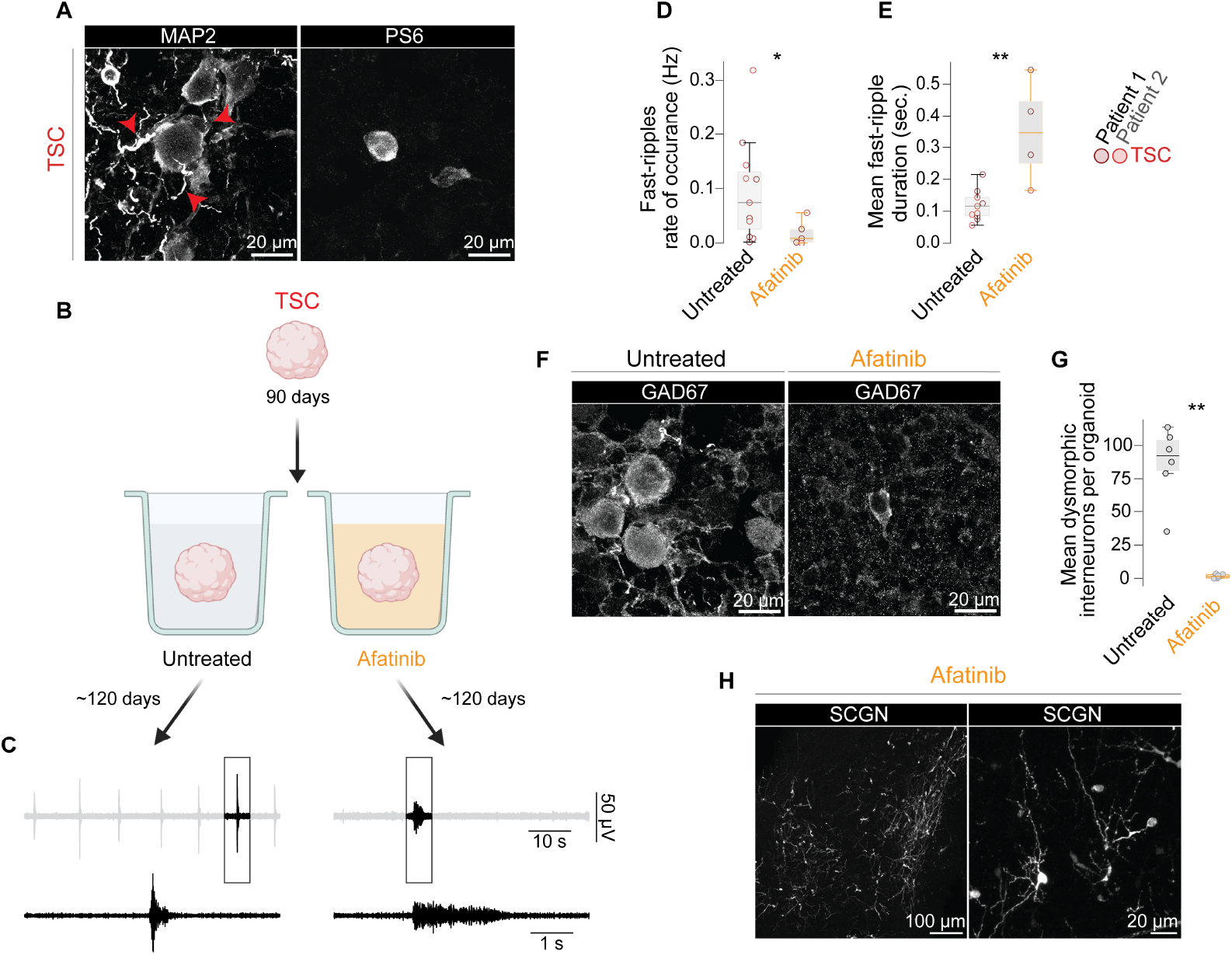
TSC organoids can uncover anti-epileptogenic drug effects. (A) Representative immunofluorescence images of dysmorphic neurons in TSC organoids show that they are neuronal (MAP2+) and have elevated mTOR signaling (PS6+) in untreated conditions. MAP2+ processes are highlighted (red arrows). (B) Schematic representation for long-term treatment of human TSC patient-derived cerebral organoids with afatinib from around 3-months (day 90). Untreated and treated organoids were matured for an average of approximately 7-months, respectively. (C) Representative fast-ripple signals from single-channels in TSC organoids before and after long-term afatinib treaments. An highlighted segment (grey box) of the original fast-ripple signal is shown with a magnified view below. (D) to (E) Quantification of the mean number of fast-ripples (untreated, n = 11; afatinib, n = 5) and their durations (untreated, n = 11; afatinib, n = 4), pre- and post long-term afatinib treatments for TSC organoids. Event durations were not calculated for organoids without detected fast-ripple events. (F) Immunofluorescence image comparisons of GAD67 in TSC organoids show that interneurons have dysmorphic morphology in untreated conditions (left), while interneurons have healthy morphology after long-term afatinib treatment (right). (G) Mean number of dysmorphic interneurons per TSC organoid in untreated (n = 3, 2 organoid sections each) and long-term afatinib treatment groups (n = 3, 2 organoid sections each). (H) Immunofluorescence images of SCGN interneurons in TSC organoids after long-term afatinib treatment highlights a reduction in dendritic pathology.

## Discussion

The lack of human models capable of investigating the neurodevelopmental origin of epilepsy poses a major challenge to developing effective therapies. Here, we use cerebral organoids derived from TSC patients to recapitulate changes in neural network activity seen in epilepsy patients and demonstrate the value of this organoid model for mechanistic analysis and evaluation of therapeutic approaches.

Epileptogenic foci in TSC are confined to cortical tubers and do not require input from other brain areas for epileptogenesis initiation ^65,66^. As cortical tubers arise during brain development ^21^, our developmental model that recapitulates tuber cell types ^18^ enabled us to assess the emergence of pathological network dynamics on a local circuit level. In this study, we show that HFOs recorded in TSC organoids closely mirrored the features observed in cortical tubers from children with TSC. These pathological events reflect the abnormal summation of synchronous action potentials from inter-connected neurons ^17,67,68^, leading to network excitability and instability ^69^. Ripples emerged more frequently with higher amplitudes ^70^, entropy, and shorter durations in TSC, further indicating abnormally enhanced neuronal synchrony and excitability ^71^. Given that single cells rarely fire faster than 400 Hz, the observed high-frequency power shifts, especially in the fast-ripple range (above 250 Hz), are likely caused by in-phase synchronous firing between neuronal ensembles ^72^. The consistent alignment of pathological ripple peaks to specific phases of the theta cycle observed in TSC organoids supports a pathologically entrained network, rather than random or desynchronized ensemble firing. The consequence of such synchronization likely explains the more frequent co-occurrences between IEDs, ripples, and fast ripples observed in TSC and other epilepsies ^68^. Additional aspects of network size and neuron distribution might also contribute to this phenotype. Notably, small clusters of neurons of dimensions that are recapitulated in organoids can generate high-frequency activity in dysplastic regions ^65^ when their firing is sufficiently synchronized ^73^ even across spatially dispersed cells ^67^. Thus, large-scale networks are not a prerequisite for generating high frequency oscillations. Taken together, our TSC organoid model enabled us to dissect local pathological network activity during epileptogenesis in cortical tubers, driven by synchronous neuronal firing.

In development, KCC2 expression regulates chloride homeostasis, shifting the depolarizing effect of GABA to a hyperpolarizing one across maturation. However, in disease conditions, abnormal changes in intracellular chloride can still cause depolarizing effects ^34^, and in epileptic tissue a similar phenomenon has been reported ^74^. Here, we show that pathological network activity in TSC organoids is accompanied by impaired maturation and a lack of periodic nested oscillations during network events, previously shown to be initiated by glutamatergic- and maintained by GABAergic signaling ^15,75^. Such changes in maturation and oscillatory events may suggest disruptions in KCC2 expression, ^17^ however, pharmacological perturbations with GABA, acting primarily through GABAA receptors, revealed clear inhibitory effects at the network level in both TSC and control organoids. While GABA could still be depolarizing in interneurons at early stages ^76^, our late 6-month time points likely precludes this ^77^. This suggests that the developmental GABA polarity switch is not affected in early epileptogenesis in TSC and such changes in epileptic tissue may rather result from chloride homeostatic alterations commonly found after recurring seizures ^74^. Together, these data suggest a model in which inhibitory GABAergic transmission plays a role in regulating pathological network dynamics.

The high temporal and spatial resolution of our silicon probe recordings allowed us to identify dysregulated firing properties of both inhibitory and excitatory neurons during pathological HFOs. In the broader field of epileptogenesis, the contribution of different neuron classes to network dysfunction remains a long-standing question. Previous stud-ies in animals have focused on dysregulation through reduced inhibition from fast-spiking parvalbumin interneurons to pyramidal neurons, and interneuron cell-death as the primary causes of cortical hyperexcitability ^78–80^. However, in this study we analyzed a confined neural network of predominantly excitatory and CGE-interneurons and found that both populations are required for the development of pathological network activity in TSC. In the context of human development, the protracted neurodevelopmental timeline and specific expansion of cortical CGE interneurons ^81–83^might indicate more diverse roles for different subtypes of GABAergic interneurons within a developing network, which is reflected in the organoids. While excessive excitation is believed to drive focal seizures, synchronized interneuron firing has been reported to occur during interictal periods before seizure onset ^84^. Our data uncover a strongly increased activity of a putative fast-firing excitatory neuron class in TSC, suggesting a network imbalance that favors hyperexcitability. However, this is accompanied by a synchronization of CGE-interneurons revealing an important role of this previously uncharacterized cell type in pathological HFO activity and network excitability. While MGE-derived interneurons primarily mediate direct inhibition, CGE-derived interneurons are more commonly associated with disinhibitory circuits ^81,82^. Therefore, a possible explanation for the role of CGE-interneurons is that they form mutually inhibiting networks, which when exposed to excitation, generate synchronized activity within the ripple and fast-ripple frequency range ^85^.

Although we previously observed increased proliferation of caudal late-born interneuron progenitors (CLIP cells) in TSC ^18^, the modest rise in activity attributed to inhibitory neurons highlight that not all CLIP-derived interneurons are functionally integrated into the network. This suggests that additional mechanisms are present, such as dysfunctional interneurons, which may underlie the impaired inhibition characteristics found in epileptic tissue ^86^. Supporting this, we discovered dysmorphic GAD67+ interneurons are only present in TSC organoids. Given their aberrant nature, we further cannot exclude the possibility that dysmorphic interneurons may release excitatory neurotransmitters in addition to GABA ^87^. All proposed mechanisms will disrupt the coordination of reciprocal excitatory-inhibitory connections that is the basis for more complex network patterns such as oscillatory activity. Accordingly, we can disentangle the alternating excitatory and inhibitory firing dynamics on single unit level in control organoids that shape complex oscillatory patterns. The disruption of these networks in TSC is reflected in the changes in single cell firing dynamics and the lack of oscillatory activities. Alterations in synaptic inputs of either excitatory neurons or interneurons can alter the timing and phase of subsequent firing in both excitation-inhibition and inhibition-inhibition loops, which may prevent oscillation outcomes as seen in TSC. Thus, in the wider context of epileptogenesis, our brain development model recapitulates a network long before seizure onset, revealing changes in both firing rate and timing of these major classes that have not been previously described in the context of pathological HFOs in epilepsy. Taken together, our model identifies a profound dysregulation in the network early during epileptogenesis involving both excitatory and inhibitory activities.

Advancing our understanding of epileptogenesis could lead to improved therapeutic concepts. Current treatments are usually initiated upon occurrence of the first seizure. However, the growing evidence for progressive network dysfunction before the first seizure has initiated a quest for preventative therapies. Such approaches are urgently needed, as the rates of drug-resistant epilepsy have shown little improvement in the last decades ^9,10^. Importantly, children with TSC at high risk for epilepsy and neurodevelopmental delay can be diagnosed before the first symptomatic seizure ^88^ based on pathological EEG alterations ^89–91^. However, despite initial hope for preventive effects in early trials with the standard anti-seizure medication Vigabatrin, a GABA transaminase inhibitor ^92,93^, a recent multicenter randomized double-blind placebo-controlled trial did not find effects on the incidence of seizures and drug-resistant epilepsy ^94^. Thus, due to the lack of appropriate model systems, the underlying mechanisms that must be targeted during epileptogenesis remain unknown and current prevention resorts to anti-seizure medications that have no anti-epileptogenic effect. Here, we use the EGF-Receptor inhibitor Afatinib to prevent epileptogenesis by targeting CLIP cell-derived interneuron production during developmental periods. After long term treatment, biomarkers for epileptogenesis such as pathological fast ripples were remodeled to reflect physiological network activities. Notably, this preventive effect occurred after prolonged EGFR inhibition during neurodevelopment. We have previously shown that Afatinib reduces proliferation of TSC tumor that contain EGFR-positive CLIP cells. Whereas it cannot be excluded that Afatinib acts on other cell types besides interneuron progenitors, EGFR expression is mainly found in the ganglionic eminences and more specifically in the caudal ganglionic eminence that also harbors CLIP cells. Furthermore, we find a specific reduction of dysmorphic interneurons and resulting downstream pathological excitotoxic features such as dendritic beading in interneurons. This suggests, that Afatinib treatment directly acts through CLIP cell-derived interneurons and that this cell type is actively involved in initiating pathological network states. Taken together, our study provides a model system to investigate the cellular mechanisms behind epileptogenesis, allowing us to demonstrate the feasibility of preventive anti-epileptogenic drug testing, further supporting the concept of early seizure and epilepsy prevention. This highlights cerebral organoids as a developmental human brain model, capable of recapitulating disease onset and progression that could bridge the gap in knowledge for better targeted therapies.

### Limitations of the Study

Certain limitations remain to be considered. Organoids recapitulate local network maturation and dysfunction, which is beneficial when modelling confined cortical tubers in TSC; however, non-focal disorders might involve coordinated activities from distal brain regions. Due to organoid limita-tions in size, complexity, and complete neural lineages, we are likely unable to reach the complete spectrum of pathological network progression. Our network-level analysis of organoid activity recapitulated electrical biomarkers relevant for clinical decision making. Nevertheless, the high resolution of the assays used in our work inherently limit the sampling of the network to individual regions and a small number of neurons. In the context of the developing human cortex, it would be highly valuable to gain a deeper understanding of HFO formation, coupling, and single-unit firing patterns from the entire network that is involved. Additionally, recordings from epilepsy patients with increased resolution could facilitate our understanding of single cell firing dynamics within and around tuber regions. Our goal was to understand the epileptogenic mechanisms during brain development, but a full characterization of this process would require longitudinal recordings across organoid development until seizure-like events occur; however, whether such activities can be recorded in brain organoids remains to be explored.

Taken together, we present a developmental brain organoid model recapitulating the focal epileptogenesis that occurs in TSC. Only by recreating and dissecting this process in vitro, will we be able to develop targeted preventive therapies that are urgently needed.

## Supporting information

Supplementary Video S1. Representative calcium imaging recording from a Control organoid.

Supplementary Video S1. Representative calcium imaging recording from a TSC organoid.

## Acknowledgments

We thank T. Lendl and the BioOptics Core Facility for help with image processing. We thank J. Sarnthein for help with signal processing. We thank O. Kim for important feedback and assistance with electrophysiology recordings. We thank C. Wilkinson for discussions on EEG analyses. We thank the Neuroscience ‘Cluster of Excellence’ (COE16) for valuable scientific discussions on CGE-derived interneurons and their role in epileptogenesis. We especially thank all patients and their families for participating in this study for donating tissue.

## Author Contributions

Conceptualization, OLE, SNW, MZ, JAK; Methodology, MZ, OLE, SNW, MAvtK, JMGV; Software, MZ, SJG, MMP, CF; Formal analysis, MZ, OLE, SNW, MavtK; Investigation: SNW, MZ, OLE, SJG, CK, SK, JC, SGG, LBM; Resources, JMGV, MFP, MaiZ, PJ, ManZ, CB; Data Curation, MAvtK, MZ, OLE, SNW, CK, MMP; Writing, OLE, SNW, MZ, JAK; Writing – Review and Editing, OLE, SNW, MZ, JAK, MAvtK, DR, RN, MaiZ, CB, PJ, NSC, JAK; Visualization, OLE, SNW, MZ, MavtK; Supervision, JAK; Funding acquisition, JAK.

## Materials and Methods

### Patient sample selection and cell line generation

Cell lines used in this study were generated in Eichmüller et al. 2022^18^. The study was approved by the local ethics committee of the Medical University of Vienna (MUV).

### Culture of induced pluripotent stem cells

iPSC culture in this study was performed as published previously ^18^. Briefly, cells were cultured in the Cellartis DEF-CS 500 culture system (Takara) according to the manufacturers’ instructions. Cells were split every third day and 400,000 cells were seeded per well of a 6 well plate. Cells were banked at different passages. For experiments, cells were used from passage 40 to 80. Genomic integrity was analyzed at passage 40 on an Infinium PsychArray v1.3 (Illumina) and compared to data from PBMCs.

### Organoid generation

Organoid generation followed a previously published method ^18^. To generate organoids, 400,000 cells were seeded into a 6-well plate. Before confluency (after 2 days), cells were washed (DPBS-/-) and incubated with 300 µl Try-pLE Express (Thermo Fisher Scientific) for 5 minutes. 1 ml supplemented Def-CS (basal medium + GF1,2,3) was added, iPSCs were triturated and transferred to a tube supplemented with 700 µl Def-CS. After counting, the required number of cells was added to a new tube. Cells were spun down (120 x g) for 3 min and resuspended in the required amount of mTeSR-1 (Stem Cell Technologies) + Rock Inhibitor (RI, 1:100): 9000 cells were seeded per well of a low attachment 96 well plate, in 150 µl mTeSR-1 + RI. Embryoid bodies (EBs) were fed 2 and 4 days after EB generation with mTeSR-1. On day 5, medium was replaced with Neural Induction (NI) medium [DMEM/F12 (Thermo Fisher Scientific), 1% N2 Supplement (produced in house), 1% MEM-NEAA (Sigma Aldrich), 1% Glutamax (Thermo Fisher Scientific)]. NI medium was changed on day 7 and day 9. On day 10, EBs were embedded in a droplet of Matrigel (Corning) and transferred to 10 cm dishes. Embedded EBs were cultured in High Nutrient Medium - Vitamin A [HN-A: 50% DMEM/F12 (Thermo Fisher Scientific), 50% Neurobasal (Thermo Fisher Scientific), 1% N2 Supplement (produced in house), 2% B27 - Vitamin A (Thermo Fisher Scientific), 1% Glutamax (Thermo Fisher Scientific), 0.5% MEM-NEAA (Sigma Aldrich,), 1% Penicillin/Streptomycin and 0.025% Insulin solution (Sigma Aldrich)]. Organoids were cultured under stationary conditions until day 13, then medium was changed to High Nutrient Medium + Vitamin A [HN+A: 50% DMEM/F12 (Thermo Fisher Scientific), 50% Neurobasal (Thermo Fisher Scientific), 1% N2 Supplement (Thermo Fisher Scientific), 2% B27 + Vitamin A (50X, Thermo Fisher Scientific), 1% Glutamax (Thermo Fisher Scientific), 0.5% MEM-NEAA (Sigma Aldrich), L-Ascorbic Acid solution (Sigma Aldrich, 7mg/ml in DMEM/F12), 1% Antimycotic/Antibiotic (Gibco), 0.025% Insulin solution (Sigma Aldrich), 1g/l Bicarbonate (Sigma Aldrich)] and organoids were moved to orbital shakers. Organoids were fed twice a week.

On day 40, organoids were transitioned gradually into Low-Nutrient medium [LN: BrainPhys Neuronal Medium (Stem Cell Technologies)3, 2% B27+A (50X, Thermo Fisher Scientific), 1% N2 supplement (Thermo Fisher Scientific), 200 nM Ascorbic Acid (Sigma Aldrich), 0.2% CD Lipid Concentrate (Thermo Fisher Scientific), 1% Antimycotic/Antibiotic (Gibco), 1% Matrigel (Corning), adjusted Glucose to 10 mM, freshly before use: 20 ng/ml BDNF (Stem Cell Technologies), 20 ng/ml GDNF (Stem Cell Technologies) and 1 mM db-cAMP (Santa Cruz)]. Gradual transfer from HN+A to LN was performed in 4 feedings (1. HN+A 75%:25% LN, 2. HN+A 50%:50% LN, 3. HN+A 25%:75% LN, 4. HN+A 0%:100% LN).

### Dorsal patterning of organoids

The dorsal patterning protocol was performed as described in Eichmüller et al. 2022^18^: organoids were patterned with HN+A medium containing 3µM CHIR99021 (Merck) from day 13 - 16.

### Silicon probe recording acquisition

Neural organoids were embedded in a block of 4% Low Melting Agarose (Promega, Cat. No. V2111) in PBS and transferred into a recording chamber (RC-42 LP, Warner Instruments) containing artificial cerebrospinal fluid (aCSF): 125 mM NaCl, 2.5 mM KCl, 1.25 mM NaH_2_PO_4_, 1 mM MgCl_2_, 2 mM CaCl_2_, 25 mM NaHCO_3_, and 10 mM D-(+)-glucose, adjusted to pH 7.4 with KOH. Embedded cerebral organoids were placed on a heated culture dish incubator (DH-35iL, Warner Instruments) and perfused with aCSF (equilibrated with a 95% O_2_ and 5% CO_2_) at a rate of ∼3–5 ml/min using a peristaltic pump (PPS2, Multichannel System). Solutions were heated by an inline heater (SH-27B, Warner Instruments) and maintained at ∼36°C with a temperature-controller (TC-344C, Warner Instruments). Acute 64 channel silicon probes with 4 shanks (ASSY-77 P-2, ASSY-77 E-1, ASSY-77 E-2, Cambridge NeuroTech) were connected to a 64-channel probe adaptor (A64 Omnetics 32×2, NeuroNexus). Signals were acquired at 30 kHz per channel and filtered at 0.1 Hz (high-pass) and 5 kHz (anti-aliasing) using the Intan 1024ch recording controller (RHD2164, Intan Technologies) through a USB 3.0 interface board (RHS2000, Intan Technologies) under the Open-Ephys plugin-GUI (https://github.com/openephys/plugin-GUI). Before electrode insertion ∼200 µm deep into the organoids, probes were immersed in a 1% (mass/vol) Tergazyme (Merk, Cat. No. Z273287) in ultrapure water (Milli-Q, IQ 7000, Merck) solution for 30 minutes at 50°C, then washed with iso-propanol and rinsed with distilled water. A minimum 5-minute baseline recording of spontaneous activity was obtained. To protect donor privacy, all organoid samples were pseudonymized prior to silicon probe recordings by assigning each sample a unique experimental identifier. The mapping key linking experimental IDs to donor identities was maintained separately.

### Intraoperative ECoG (ioECoG) recordings

Two children with a proven *TSC1* mutation were selected from the cohort of participants of the HFO trial, a randomized controlled trial comparing the use of spikes and HFOs for intraoperative neurosurgical guidance, in whom long term 4-year post-surgical seizure freedom (ILAE class 1) was achieved. The medical committee at UMC Utrecht approved the trial protocol (MEC-13389). The results of the HFO trial were published separately ^29^. Intraoperative electrocorticography was recorded using a 4×5 electrode grid with (Pat#2) or without (Pat#1) a 1×8 electrode strip (Ad-Tech, Racine, WI, USA) placed directly on the cortex. Recordings were acquired with a 64-channel (MicroMed LTM Express, MicroMed, Veneto, Italy) or 32-channel (MicroMed Flexi, MicroMed, Veneto, Italy) EEG-system at a 2048 Hz sampling rate with an antialiasing filter at 538 Hz and high pass filter at 0.15 Hz. The signal was referenced to an external electrode placed on the mastoid. Propofol anesthesia was temporarily paused during the ioECoG recordings to avoid burst-suppression patterns and suppression of epileptic activity in ioECoG recordings. Once a continuous background pattern was reached, signals were recorded for at least 2 minutes before moving the electrode grid to the next position. As both children were allocated to the HFO arm, the ioECoG was reviewed intraoperatively for HFOs in Stellate Harmonie Viewer (Natus Medical Inc., Montreal, QC, Canada) to guide the resection. Intraoperative photographs of the ioECoG grid positions on the cortex were taken ^29^.

### Clinical outcome after HFO based ioECoG-tailored surgery

In the first patient (Pat#1) pathological interictal activity was recorded over two lesions, including frequent ripples with co-occurring fast ripples and a continuous peak pattern. Both lesions were resected, resulting in absence of epileptiform activity over cortical tissue directly surrounding the resection cavities. Notably, although complete removal of all pathological HFOs was not possible as the resection was limited by function in the central (sensori-)motor region, the lesionectomy resulted in seizure-freedom (Figure 2A and Table S1). In a second patient (Pat#2), 3D magnetic resonance imaging (MRI) visualized two cortical tubers in the right temporal lobe (Figure S2A). ioECoG recordings revealed interictal pathological activity over these tuber regions, including pathological ripples and fast ripples (Figure S2B). First the posterior lesion was resected and in a second step the anterior lesion, thereafter no epileptogenic ripples or fast ripples remained, allowing sparing of the mesiotemporal regions while achieving post-surgical seizure-freedom (Table S1).

### Multi-unit spike detection and analysis

To detect multi-unit spiking activity, extracellular recordings sampled at 30,000 Hz were band-pass filtered between 300 and 3,000 Hz using a forward-backward-zero-phase 4th-order Butterworth filter. Noisy channels were automatically removed with SpikeInterface ^56^ (version 0.101.0, https://github.com/SpikeInterface) using both the coherence between channels and the power characteristics of individual channels. The resulting filtered signals were then re-referenced to the median signal (common median reference). Negative peaks were detected in each channel independently above a threshold 5 times the median absolute deviation (MAD) using a `peak_detection` function ported from Tridesclous into SpikeInterface. Multi-unit spikes were abolished with tetrodotoxin applications (Figures 1E, S1E and G), confirming robust detection methods. Mean firing rate calculations were conducted across ‘active’ channels, defined as those with a firing rate > 0.1 Hz

### Network event analysis

Detected multi-unit spikes were converted into binary vectors and summed across all available channels (up to 64) within 1 ms window bin sizes and subsequently smoothed with a 25 ms kernel. Putative peaks of network events were identi-fied using the resulting network event array, provided they exceeded at least twice the standard deviation (2×STD) above the noise floor. Start and end times were defined by time windows surrounding the peaks of the network events, when the amplitude was reduced to 0.2 times the standard deviation (0.2×STD). Network events start times less than 500 ms apart were automatically merged and removed if at edges. Resulting network events were visually inspected and manually curated. SD thresholds or amplitude cut-offs were adjusted to account for lower activity recordings. To be included for subsequent analyses, a network event was required to meet the following criteria: i) at least 0.5 spikes/ms; ii) a duration between 10 ms and 10 seconds; and iii) 1 second away from adjacent events. Network events were smoothed with a Savitzky-Golay filter (100 ms window length, 1st order polynomial) for display purposes (Figures 1D and 1I), but not for subsequent analyses. Network events of varying durations were standardized by padding with zeros for display purposes only (Figure 1I). Analysis was performed on base-line recordings across the ‘most’ active epoch, determined by the number of network events.

For network event peak detections, network events were first smoothed with a Savitzky-Golay filter (100 ms window length, 5th order polynomial). The resultant signal was then normalized to the max peak. Subpeaks were detected after the max peak of a network event if they had an ii) normalized absolute height exceeding 10%, ii) peak prominence greater than one-half of the absolute peak height, iii) duration > 25 ms, and were iiii) at least 50 ms apart from adjacent peaks. Peak prominence values were adjusted to at least 5% (0.05) for recordings with low active channel numbers. All peak isolations were cross-examined against the underlying local field potential (LFP) voltage changes and mean network event kernels (Figures 1D and S1I).

### High frequency oscillation (HFO) detections

Silicon probe extracellular recordings (sampled at 30,000 Hz) were filtered using a forward-backward-zero-phase finite impulse response (FIR) filter (10 Hz transition band), with cut-off frequencies set between 80 and 250 Hz for ripples and 250 and 500 Hz for fast ripples. Filtered data was then down-sampled to 2,000 Hz. In organoids, HFOs were detected in ripple and fast ripple signals using their amplitude envelope, which was derived using a Hilbert transform and further smoothed with a 20 ms sliding kernel. Start and end times were identified when the standard deviation (STD) of the root-mean-square (RMS), calculated over 5 ms segments, exceeded 5 STD and crossed 3 STD. SD thresholds were adjusted where necessary for individual recordings. HFOs were included in subsequent analysis if their durations ranged between 20 ms and 600 ms, consistent with previous reports. Events less than 10 ms apart were merged. All events were visually inspected and manually curated (Figure S3C), in accordance with HFO detection consensus guidelines ^46,95^. Detected HFOs had morphologies consistent with previous reports ^46^. All detections and subsequent analyses were conducted on the channel with the highest amplitude, determined by maximum ripple amplitudes. Recordings without detected HFOs were excluded from analyses to avoid biases.

For ioECoG recordings, signals preceding the first re-section were labeled as “pre-ioECoG,” whereas the recordings after the final resection were designated “post-ECoG.” Intermediate recordings were excluded from further analysis. One-minute epochs at the end of each ECoG recording were selected for subsequent analysis to minimize propofol effects ^96^. For HFO detections in the ioECoG data, an automated Matlab-based HFO detector was used with a zero-phase FIR filter applied between 80-1000 Hz for ripples, and 250-1000 Hz for fast ripples ^95,97^. To optimize HFO threshold settings for the ioECoG data, a training set consisting of four patients was used. The output of the HFO detector included the start and end time on the channel of occurrence per detected HFO ^98^. All detected HFOs were visually checked to correct for artifacts as well as false-positive and missed HFOs by MvtK using Brain Quick Software (Micromed SpA). Visual classification criteria for physiological ripples were presumed eloquent cortex, no IED co-occurrence, and lower amplitude and/or longer duration ^70^. A split dual screen was used to simultaneously visualize ripples (FIR filter, >80 Hz; gain, 50 uV/mm, 1s/page) and FRs (FIR filter, >250 Hz; gain, 10 uV/mm. 1s/page) for visual inspections and manual curations. Channels with the highest rate of pathological or physiological ripple occurrence were selected for subsequent analyses.

For display purposes and signal feature extractions, both ioECoG and silicon probe recordings were filtered between 80 and 250 Hz for ripples, and 250 and 500 Hz for fast ripples using a forward-backward-zero-phase FIR filter from the python package, MNE (version 1.6.0, https://github.com/mne-tools/mne-python). HFO amplitudes were calculated from the Hilbert transform of the filtered signal between identified and curated start and end times. Entropy was calculated using an approximate entropy algorithm with the antropy package (version 0.1.6, https://github.com/raphaelvallat/antropy). The Chebyshev distance metric was applied during this calculation. Differences in durations were accounted for by normalizing approximate entropy to the number of samples. Epochs of highest activity were isolated for feature extractions in organoids. For the ioECoG data only the last one-minute per recording situation was analyzed to avoid propofol effects.

### Interictal epileptic discharge (IED) detection

Extracellular signal recordings were bandpass filtered with cutoff frequencies between 20 and 80 Hz using a forward-backward-zero-phase FIR filter (2 Hz transition band). Silicon probe recordings (sampled at 30000 Hz) were then down-sampled to 2000 Hz. Electrodes chosen for IED detections in organoids were matched to highest amplitude channels previously identified in high frequency oscillation (HFO) detections. Powerline noise (50 Hz) was excluded from subsequent analysis. IEDs were detected on the amplitude envelope, which was derived using a Hilbert transform and further smoothed with a 10 ms sliding kernel. Peak times were identified when the standard deviation (STD) of the root-mean-square (RMS), calculated over 5 ms segments, exceeded 10 times the baseline STD. IED start and end times were marked when the signal crossed 5 times the baseline STD. For organoids, IEDs were included if the following criteria were met: i) durations between 10 and 100 ms, ii) occurring greater than 200 ms from another with iii) amplitudes greater than 10 µV, calculated using the smoothed amplitude envelope. All events were then visually inspected for accuracy. Events were manually removed if artifacts were present (Figure S3C). Detected IEDs in organoids had waveforms consistent with typical interictal spikes (Figures 3B and H) ^51^. For human recordings, IEDs were detected as sharp transient deflections in the IED signal (Figure 2C) above the baseline, accompanied by sharp spikes in the LFP recordings. IEDs were confirmed visually and displayed morphologies consistent with previous reports ^51^.

### Pathological high frequency oscillation classifications

In cerebral organoids, pathological HFOs (pHFOs) were classified if detected IEDs (80 - 250 Hz), ripples (80 - 250 Hz) and fast-ripples (250 - 500 Hz) were found to occur simultaneously within a 200 ms epoch. Given that pathological HFOs in intraoperative EcoG recordings from children with TSC occur on sharp transient spikes in corresponding LFP signals (Figure 2B), we further detected sharp peaks in LFP (0.5 - 1000 Hz) signals beyond an absolute voltage of 30 µV and a 5 seconds horizontal distance from neighboring events. Peaks with a prominence above 5 µV were kept. Resulting events were visually inspected and manually curated to confirm morphology similar to patient pHFOs. Accordingly, all pHFOs coincided with sharp transients in the LFP signal (Figure 3B and H).

### Spike sorting

Single unit spike sorting, pre-processing and post-processing processes were performed using the unified framework, SpikeInterface ^56^ (version 0.101.0). Spikes were assigned to individual neurons as follows: raw extracellular recordings were bandpass filtered 300-3000 Hz using a forward-backward zero-phase 4th order Butterworth filter. Noisy channels were identified and automatically removed following a procedure used in *multi-unit spike detection and analysis*. Each channel was subsequently re-referenced to the median signal (common median reference). The Kilosort2 algorithm was then applied to the filtered signals to extract single-unit activity within a channel radius sparsity of 200 µm, using a detection threshold of 5 rms above the baseline and a minimum firing rate of 1/600 Hz. Spike times in a spike train that occurred within a 0.3 ms period were considered duplicate and subsequently removed. Kilosort2’s results were then subject to both manual and auto-curation techniques to ensure the physiological relevance of each single-unit.

Quality metrics were automatically calculated for each single unit based on their waveform, firing rate and inter-spike interval (ISI) distribution using the `compute_quality_metrics` function ^56^. Firstly, units that had a signal-to-noise ratio (SNR) of > 2 and firing rate > 0.05 Hz were manually curated using the Phy GUI (version 1.0.9, https://github.com/cortex-lab/phy). This process involved evaluating the autocorrelograms of single units, along with the spatial distribution and morphology of their waveforms. Units that followed the biophysical properties of an action potential generation were included, such as coherent waveform patterns across neighboring channels. The cross-correlograms, mean waveforms and waveform spatial distributions of single units with high similarities (> 0.8) were checked and merged if considered similar. In several cases, spike units were split if distinct waveform footprints were seen across Phy’s waveform and feature space viewers. Positive-spiking units were manually excluded, due to their association with dendrites and axons.

An additional auto-curation process was then applied to spike units following manual curations. This ensured that spike-units in the final dataset had an (i) ISI violation rate of ≤ 1%, (ii) firing rate *>* 0.1 Hz, (iii) presence ratio *>* 0.8, and (iv) signal-to-noise ratio ≥ 2. The ISI violation rate was calculated for each unit as the total number of ISI violations, divided by the total number of spikes. The threshold for identifying refractory period violations was set to 1.5 ms. Post curations, spike features were calculated using the templates of detected units with SpikeInterface’s template_metrics function. Spike-unit waveforms were extracted across all active channels within a 3 ms (90 samples) window, averaging up to 500 spikes. Asymmetry was additionally calculated using maximum (negative) amplitude mean waveforms, normalized to the spike minimum (Figure S5B).

### Spike-unit waveform clustering

To classify neuron-types, an unsupervised classification was performed using single-unit waveforms isolated in spike sort-ing. Putative cell-type clustering was performed using the WaveMAP Python package ^59^, which combines non-linear dimensionality reduction (Universal Manifold Approximation and Projection [UMAP]) with Louvain clustering methodologies. To control for the impact of morphological structures (soma, dendrites, axons) on extracellular waveform shapes, we isolated the largest negative amplitude waveform for each unit. Large negatively deflecting spikes are reported to mostly reside at the soma. Before clustering, single-unit max waveforms were normalized between -1 and +1 by first subtracting their baseline (calculated as the first 20 samples) and then scaling by the value of its most negative amplitude. Hyperparameters for UMAP embeddings were configured with 20 neighbors, a minimum distance of 0.1, and a random state of 42. The Louvain clustering resolution was set to 2, as previously described. For visualization only, waveforms were shown in figure 4B with a Gaussian filter smoothing applied (using a standard deviation of 2), and an amplitude-preserving scaling. A gradient boosted decision tree classifier (xgboost version 1.7.4) was trained on a 30-70test-train split of normalized extracellular waveforms to predict classified neuron cluster types (Figure S4D). Hyperparameter tuning was performed via a grid-search and 5-fold cross-validation.

### Perievent firing rates and spike probabilities

To evaluate neuron firing patterns during HFO events (Figure 4I), spike times were aligned to the peak of detected events, as described in high frequency oscillation (HFO) detections. A peri-event window spanning 500 ms before to 2.5 seconds after the HFO peak (t = 0) was used for analysis. Firing rates were computed for each unit using 10 ms bins and averaged across all detected events (Figure 4H). Spike firing probability was determined by calculating the proportion of spikes occurring +/-200 ms from the HFO peak.

### Local field potential signal processing

Local field potential (LFP) signals were obtained by band-pass filtering the raw signals between 0.5 and 1000 Hz using a forward-backward-zero-phase 4th order butterworth filter. For silicon probe extracellular recordings (sampled at 30,000 Hz), LFP signals were down sampled to 2000 Hz. Potential powerline noise was removed using a zero-phase IIR notch filter at 50 Hz in silicon probe recordings. Time-frequency spectrograms were generated on LFP signals using a Stockwell Transform with a Gaussian window, implemented via the Stockwell package (version 1.1.2, https://github.com/claudiodsf/stockwell/tree/main). The frequency ranges of interest were 1-20 Hz for low-frequency signals, 20-80 Hz for interictal epileptic discharges, and 80-500 Hz for both ripple and fast ripple canonical bands. The power spectral density (PSD) was computed using a Fast Fourier Transform (FFT) on the LFP.

### Phase amplitude coupling analysis

Phase amplitude coupling (PAC) was evaluated between the phase of theta and the hilbert envelope of ripple signals. LFP signals were bandpass filtered between 4 and 8 Hz using a forward-backward-zero-phase FIR filter (0.5 Hz transition band). Theta phases were extracted by taking the phase angle of the theta signal Hilbert transform. Modulation index (MI) was quantified by firstly binning theta phases at 18-degree angles (20 bins) between -π and π. Ripple amplitudes were then binned by corresponding theta phases values and normalized by the sum. The MI was calculated as the Kullback-Leibler (KL) divergence between the normalized phase-amplitude distributions across phase bins and a uniform distribution. A Savitzky-Golay filter (100 ms window, 5th order polynomial) was applied to the Hilbert envelope before binning. In organoids, MI quantifications were performed across ripple-event and non-event epochs defined on highest amplitude channels. Ripple-event start times were 250 ms before a ripple peak, as identified in high frequency oscillation (HFO) detections. For each silicon probe recording, ripple peak times atleast 5-seconds apart were selected, and the end ripple-event time was set from the ripple peak as the maximum network event duration as described in network event analysis. Non-event epoch times were defined as times between ripple-event epochs. MI values were averaged across all epochs.

### Pharmacology

Pharmacological applications were applied to neural organoids in aCSF and perfused into the recording bath for at least five minutes after stable baseline recordings. CNQX (6-Cyano-7-nitroquinoxaline-2,3-dione, 10 µM, Tocris, Cat. No. 1045) blocked AMPA receptors. GABA (γ-Aminobutyric acid, 20 µM, Sigma-Aldrich, Cat. No. A2129) acted as a GABA receptor agonist. Carbenoxolone (CBX, 200 µM, Sigma-Aldrich, Cat. No. C4790) inhibited gap junction communication by blocking connexin channels. TTX (Tetrodotoxin citrate, 0.5 µM, Tocris, Cat. No. 1069) blocked sodium channels and was applied at the end of the recording to confirm neural responses. All drugs were added from frozen aliquots immediately before each experiment. To account for equilibration times, the first 5 minutes of extracellular recordings following the application of TTX, CBX, CNQX, and GABA to the recording bath were disregarded, and the subsequent 2-5 minutes were analyzed.

For long term treatment of organoids with Afatinib (Selleckchem, Cat. No. S1011), a final concentration of 1 µM was freshly added to BrainPhys media with every feeding from day 90 on. Organoids were fed twice a week until the recording day, then subsequently fixed and analyzed. When performing extracellular recordings, organoids were first recorded in aCSF supplemented with 1 µM Afatinib, then Afatinib was removed after 15 minutes to analyze with-drawal effects.

### Transmission Electron Microscopy

For electron microscopy studies, organoids were fixed with 3.5% glutaraldehyde. Samples were then post-fixed with 2% osmium, rinsed, dehydrated and embedded in Durcupan resin (Fluka). Semithin sections (1.5 µm) were cut with a diamond knife and stained lightly with 1% toluidine blue. Ultra-thin (70 nm) sections were cut with a diamond knife, stained with lead citrate (Reynolds solution) and examined under a transmission electron microscope (FEI Tecnai G2 Spirit BioTwin). Images were acquired using Radius software (Version 2.1) with a XAROSA digital camera (EMSIS GmbH, Münster, Germany).

### Immunohistochemistry on organoid samples

Organoids were fixed in 4% paraformaldehyde for 1 h at room temperature, then washed with PBS three times (3×10 min) at room temperature and then allowed to equilibrate in 30% sucrose at 4 °C. Organoids were embedded in OCT (optimal cutting temperature compound; Scigen, Cat. No. 4586) and sectioned at 20 µm on a cryostat (Leica NX70). Sections were blocked and permeabilized in blocking solution [0.25% TritonX-100 with 4% normal donkey serum (Millipore) in PBS for 1 hour at room temperature. Sections were then incubated overnight at 4°C with primary antibodies in staining solution (0.1% Triton X-100 with 4% normal donkey serum (Millipore) in PBS]. Sections were washed three times in PBS. Secondary antibodies were incubated for 2 h at room temperature and afterwards sections were washed again three times in PBS. DAPI was added to secondary antibodies to mark nuclei. Secondary antibodies labeled with Alexa fluor 488, 568, or 647 (Invitrogen) were used for detection. Slides were mounted using fluorescence mounting medium (Dako) and dried overnight before imaging.

### Microscopy

Images were acquired on LSM880 and LSM800 confocal laser scanning microscopes (Zeiss). Large overview images were acquired using the Pannoramic Slide Scanner 250 Flash II or III system (3DHistech). To image cleared organoids for beading quantifications and for live-imaging of GCaMP6s calcium transients, images were acquired using an Olympus Spinning Disk system based on the Olympus IX3 Series (IX83) inverted microscope with a dual-camera Yokogawa W1 spinning disk. A 40x/0.75 (Air) WD 0.5mm objective was used, with an additional ×3.2 magnification of the Orca Flash 4.0 camera for high magnification.

### Antibodies used

The following antibodies were used to perform immunostainings on organoid tissue: CAMKII (50049; Cell Signaling; 1/200), SATB2 (ab51502; Abcam; 1/200),CALB2 (CR6B3; Swant; 1/1000), PVALB (AF5058; RD; 1:200), SST (MAB354; Sigma-Aldrich; 1/200), GABA (A0310; Sigma-Aldrich; 1/200), PS6 (14236; Cell Signaling; 1/300), GAD67 (ab80589; Abcam; 1/300),SCGN antibody (HPA006641-100UL; Sigma-Aldrich; 1/1000 2D; 1/200 3D), MAP2 (PA1-10005; Invitrogen; 1/1000 2D; 1/200 3D).

The following antibodies were used on human brain samples: COUP-TFII (PP-H7147-00; Ms; Perseus Proteomics), Sp8 (sc-104661; Gt; Santa Cruz Biotechnology), SCGN (HPA006641; Rb; Sigma-Aldrich). All in 1:200 dilution.

### Tissue clearing and 3D immunohistochemistry

The 2Eci tissue clearing was performed as previously described. In brief, organoids were fixed in 4% paraformaldehyde for 1 h at room temperature. For 3D immunohistochemistry, PBS-TxDB solution was used (sterile filtered solution of 10x PBS, 2% TX100, 20% DMSO, 5% BSA (all per-cent volume with double distilled H2O was added to reach the final volume). Organoids were permeabilized on a rotor for 1 day in 10 ml PBS-TxDB at room temperature. Up to 3 organoids were then transferred into 2 ml Eppendorf tubes containing primary antibody solution in PBS-TxDB (between 500 µl and 1 ml) and transferred to a rotor for 5 days. Organoids were then washed in 10 ml PBS-TxDB for 2 days (3× PBS-TxDB exchange) and the same steps were repeated for secondary antibodies. After secondary antibodies were washed off, the organoids were washed in PBS for 1 h and then fixed in 4% paraformaldehyde for 1 h. Organoids were then dehydrated in a 1-propanol gradient (30%, 50%, 70%, 100%, 100%, 8 h per step) with Milli-Q water. After dehydration, organoids were transferred into ethyl cinnamate and were imaged as soon as they were completely transparent (1–3 h or overnight at room temperature).

### Dendritic beading morphology quantification

The Olympus Spinning Disk system was used to acquire confocal Z-stack images at 128x magnification. To obtain high resolution images for morphology reconstructions and quantification, each Z-stack was composed of 161 images with 0.25um Z-steps. N software was used for processing. All images were loaded into SNT plugin version 4.2.1 and each neuronal process was traced as a single path. All processes were exported, then each bead was traced manually in Fiji at the largest diameter (`diameter_big`) and the corresponding upstream process (`diameter_small`), to obtain `process_length`, `diameter_big`, and `diameter_small`. The ratio for each bead diameter to process diameter was calculated to obtain the bead size normalized to process. The mean of the ratio per process was calculated between all mutant and control neurons and plotted with R.

### Dysmorphic neuron quantification

Large overview images of TSC organoid sections with or without long-term Afatinib treatment were acquired using the Panoramic Slide Scanner 250 Flash II or III system (3DHis-tech). Quantifications were performed on two separate 20 µm organoid sections from three organoids each of the Afatinib-treated or untreated conditions. Overviews were loaded into Fiji and the Cell Counter plugin was used to track counts per section. Cells were counted as dysmorphic based on GAD67/PS6 staining’s with clear enlarged morphology. Two examiners quantified the number of dysmorphic neurons separately and the average number per section was taken.

### Immunohistochemistry on frozen human brain samples

Tissue was collected with consent in strict observance of the legal and institutional ethical regulations of the University of California San Francisco Committee on Human Research. Protocols for use of surgical tissues were obtained via the UCSF Brain Tumor Research Center. The ethics approval number for the use of de-identified human biospecimens is 10-01318. These studies were in accordance with the ethical standards of the institutional research committee and with the 1964 Helsinki declaration and its later amendments. Tissues were fixed in 4% paraformaldehyde for 2 days, and then cryoprotected in a 30% sucrose solution. Blocks were cut into 30-micron sections on a cryostat and mounted on glass slides for immunohistochemistry. Frozen slides were allowed to equilibrate to room temperature for 3 hours. 10 minutes antigen retrieval was conducted at 95°C in 10 mM Na Citrate buffer, pH=6.0. Following antigen retrieval, slides were washed with TNT (0.05% TX100 in PBS) for 10 minutes, placed in 1% H2O2 in PBS for 90 minutes, and then blocked with TNB solution (0.1 M Tris-HCl, pH 7.5, 0.15M NaCl, 0.5% blocking reagent from PerkinElmer) for 1 hour. Slides were incubated in primary antibodies overnight at 4°C and in biotinylated secondary antibodies (Jackson Immunoresearch Laboratories) for 2.5 hours at room temperature. All antibodies were diluted in TNB solution from PerkinElmer. Sections were then incubated for 30 min in streptavidin horseradish peroxidase that was diluted (1:200) with TNB. Tyramide signal amplification (Perkin-Elmer) was used for some antigens. Sections were incubated in tyramide-conjugated fluorophores at the following dilutions: Fluorescein, 1:50 (4.5 min); Opal 570, 1:100 (10 min); Cy5, 1:50 (4.5 min). Sections were acquired on a Leica TCS SP8 confocal microscope. Large overview images were imaged using a Keyence microscope.

### Calcium imaging recordings and analysis

TSC patient and control iPSC lines were genetically modified to express the calcium indicator GCaMP6s under the human synapsin promoter (TSCpatient1-hSyn1-GCAMP6s) using CRISPR/Cas9 genome editing in-house. Organoids were generated using the protocol detailed above and imaged at approximately 180 days in vitro (see microscopy section). Calcium transients were recorded using a 20X objective at a frame rate of 15.4 frames/second. To protect donor privacy, like for silicon probe recording acquisition, all organoid samples were also pseudonymized prior to recording. All recordings were performed in BrainPhys neuronal media at ∼37°C (equilibrated with 95% O_2_ and 5% CO_2_) and are 5 minutes and 25 seconds long. For processing, recordings were downsampled to 512 x 512 pixels and the first 100 frames were removed. Fluorescence transients were extracted using the open-source software package CaImAn (https://github.com/flatironinstitute/CaImAn) ^99^. All further analysis was performed in Python using custom codes. Extracted neuronal traces were then filtered to remove artifacts and were subsequently z-scored. Recordings with less than 3 functional neurons were excluded from the analysis. Net-work events were calculated as the sum of “global” peaks detected on the average neuronal matrix of one recording as >25% of the maximum amplitude, calculated on the average matrix. Inter-burst intervals were defined as the distance between two “global” peaks of the average matrix of neurons. The mean inter-burst interval was calculated as the median of all inter-burst intervals of the average matrix of one recording. A counter was calculated as the sum of all individual peaks that fell into a window defined around each global event. One individual peak can only be counted once. The window around each global event is defined as the average peak duration for the recording. The average peak duration was calculated as the average peak width of all “global” peaks of the average matrix of neurons. For a given batch and time point, only recordings with a corresponding control or mutant pair were included, resulting in a total of 324 recordings used for analysis. Data analysis was performed blind to experimental conditions and no data points were excluded from analysis.

### Quantification and statistical analysis

Data are presented as mean ± SEM throughout text and legends, or median with the interquartile ranges specified (e.g. boxplots). Figure legends specify n-values and P values for all statistical tests, where each n value represents an independent experiment. As biological datasets rarely follow a normal distribution, statistical analyses were performed using the two-sided Mann-Whitney U test or the two-sided Wilcoxon signed-rank test, implemented in Python using the scipy.stats package. Symbols with errors depict mean ± SEM. Kolmogorov-Smirnov tests were applied to pooled distributions, such as firing rate probability profiles, to assess differences in their cumulative distributions between experimental groups and putative neuron type. Organoid data points (unpatterned protocol) from baseline recordings were excluded from analyses if network event frequencies were greater than 3-times the standard deviation of their respective experimental groups. Sample sizes were estimated empirically based on previous studies.

### Data and code availability

Original data and analysis programs were stored in the scientific repositories of the Institute of Molecular Biotechnology of the Austrian Academy of Sciences (IMBA). All data reported will be made available upon reasonable request from the corresponding author. Custom code for data analysis and figure generation will be made available upon reasonable request from the corresponding author.

**Fig. S1.**
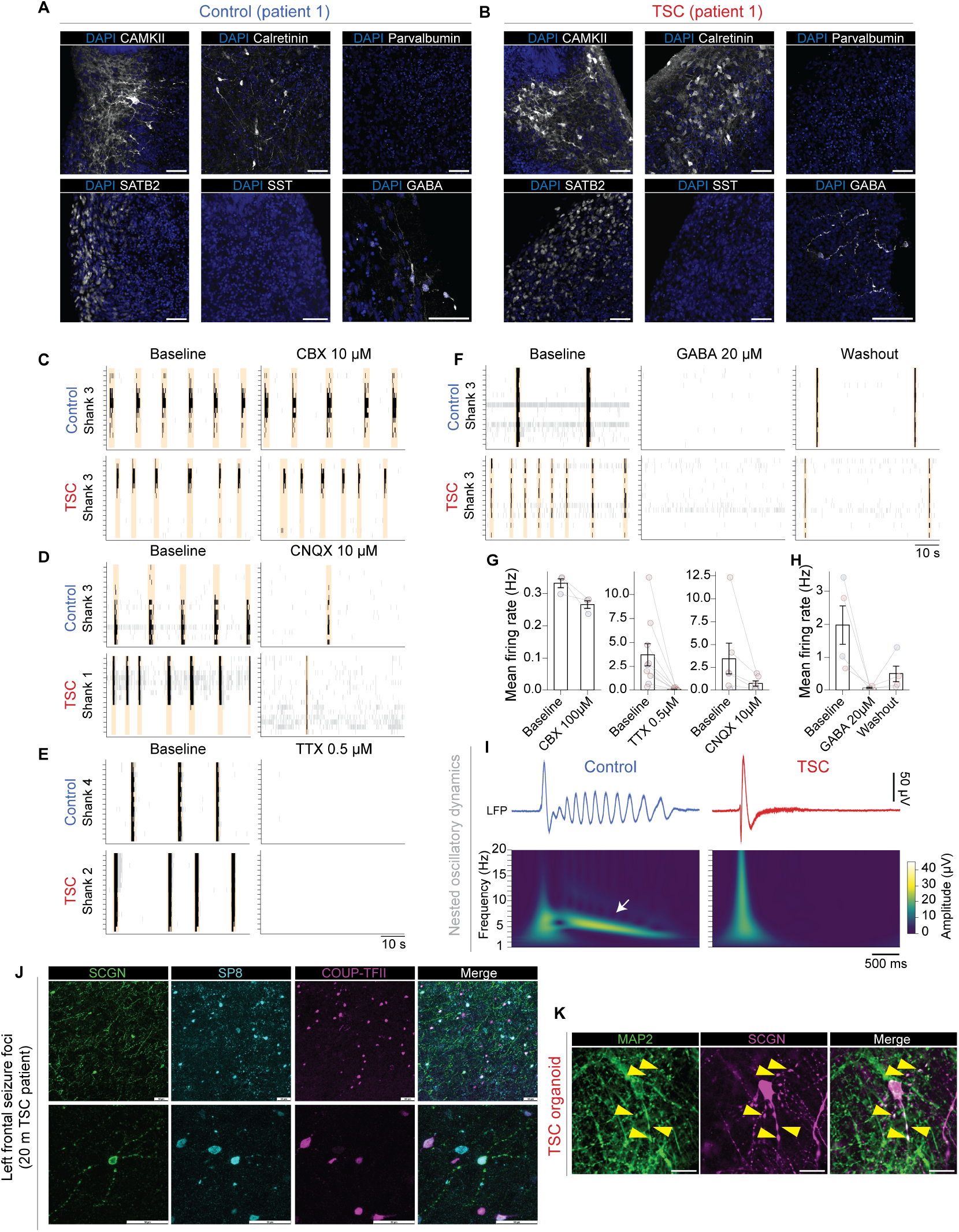
Characterization of functional neural networks in TSC, related to Figure 1. (A and B), Immunostainings on isogenic control organoids (A) and TSC patient organoids (B) from patient 1 at day 126 show excitatory neurons marked by TBR1, CAMKII and SATB2. Organoids are positive for COUP-TFII and Calretinin, and negative for NKX2.1, SST and Parvalbumin, marking GABAergic inhibitory interneurons from the CGE and not the medial ganglionic eminence (MGE). DAPI+ nuclei are shown in blue. Scale bars: 50 µm. (C to H) Pharmacological perturbations of network activity affect event occurrences and MUA firing rates in control and TSC organoids. Representative raster plots showing that blocking transmission through gap junctions with CBX (200 µM) did not affect synchronous network activity (C) and MUA firing rates (G). In contrast, CNQX (10 µM) application blocking AMPA receptors (D) and 0.5 µM TTX (E) significantly reduces synchronous network event activity and firing rates (G). Application of 20 µM of GABA to TSC and control brain organoids abolishes synchronous network activity (F) and firing activity (H). Partial recovery during washout validates the role of GABAergic transmission. Network events are highlighted in orange. (I) Top: Representative LFP signals from a single electrode channel during network events in control (n = 31 network events) and TSC (n = 24 network events) organoids between 6-7 months, presented mean ± SEM. Bottom: Corresponding time-frequency spectrograms of mean LFP signals highlight dominant power in the delta and theta range (white arrow), especially related to trailing oscillatory subpeaks. (J) (Top and bottom left) Immunohistochemistry overview of SCGN from two samples from the left front seizure focus of a 20-month TSC patient. (Top right panels) 20x magnification of the first sample shows SCGN+ interneurons within the front seizure focus that co-localize with Sp8 and COUP-TFII, confirming CGE identity. (Bottom right panels) 63x magnification of the second sample highlights one SCGN+ interneuron within the frontal seizure focus that exhibits pathological dendritic beading. This interneuron also co-localizes with Sp8 and COUP-TFII, confirming CGE identity. Scale bar: A: 50 µm; B: 50µm (K) In organoids, immunohistochemistry of SCGN in TSC organoids show beading morphology in the SCGN+ interneurons and co-localization with MAP2, marking dendrites. Yellow arrowheads mark dendritic beads that SCGN+/MAP2+.

**Fig. S2.**
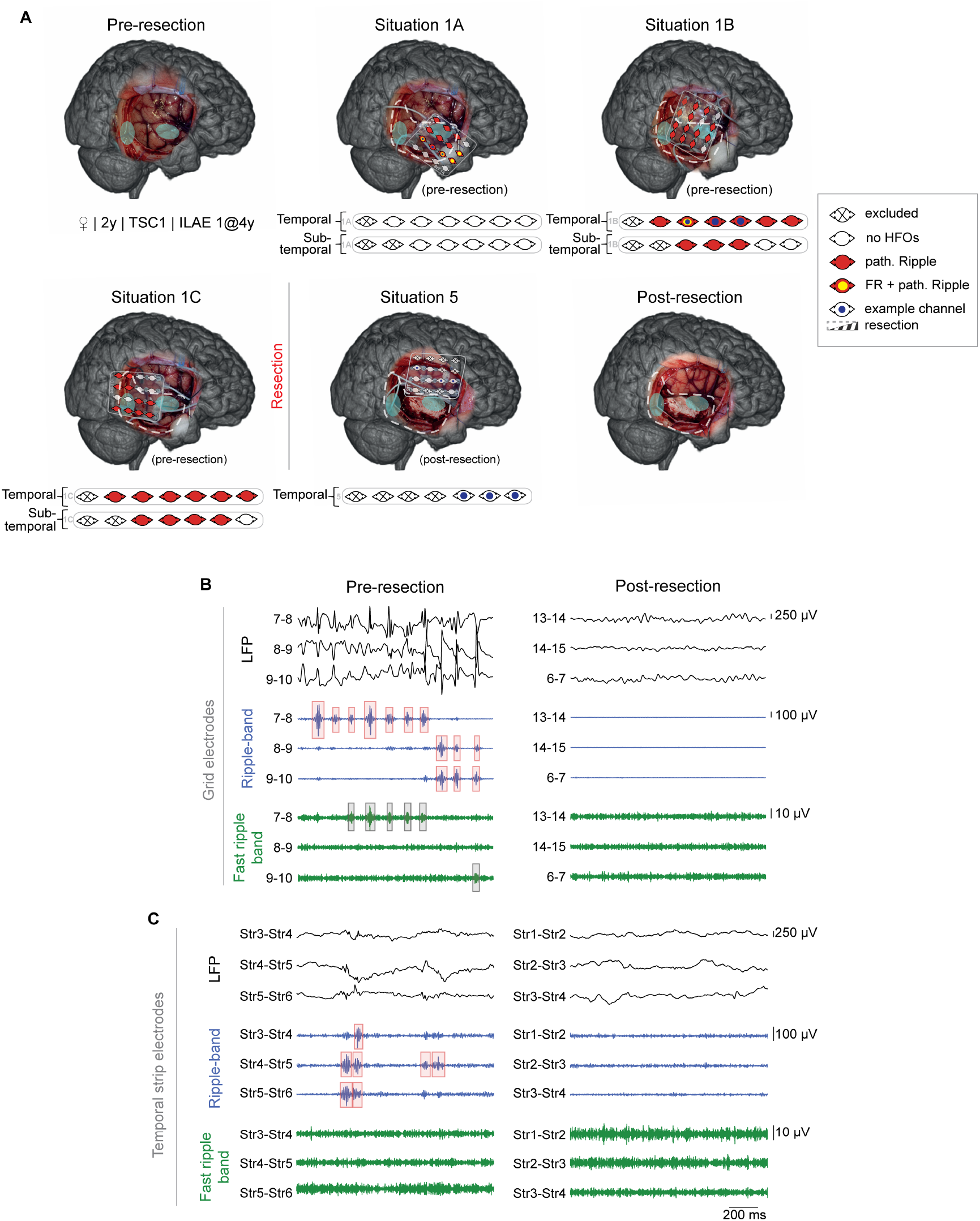
Complete removal of epileptogenic HFOs renders child with TSC seizure free, related to Figure 2. (A), Intraoperative electrocorticography recording pre- and post-resection superimposed on 3D MRI reconstruction. All pre- and post-ioECoG grid electrode positions are shown, as well as a schematic representation of the strip electrodes that were either placed under the scalp towards the temporal pole and mesiotemporal structures. The color code marks detection of pathological ripples and fast ripples, as well as the example channels shown in panels B and C. Abundant pathological ripples and fast ripples were recorded pre-resection surrounding both tubers, however, no physiological HFOs were recorded. Post-resection, no pathological HFOs were recorded. (B and C), Example of pre- and post-resection recordings in LFP, ripple-(80-250 Hz; FIR filter), and fast-ripple-band (250-500 Hz; FIR filter) recorded with grid (B) and temporal electrodes (C) marked in a. Three representative channel recordings are shown across two separate situations. Pathological ripples are annotated in red.

**Fig. S3.**
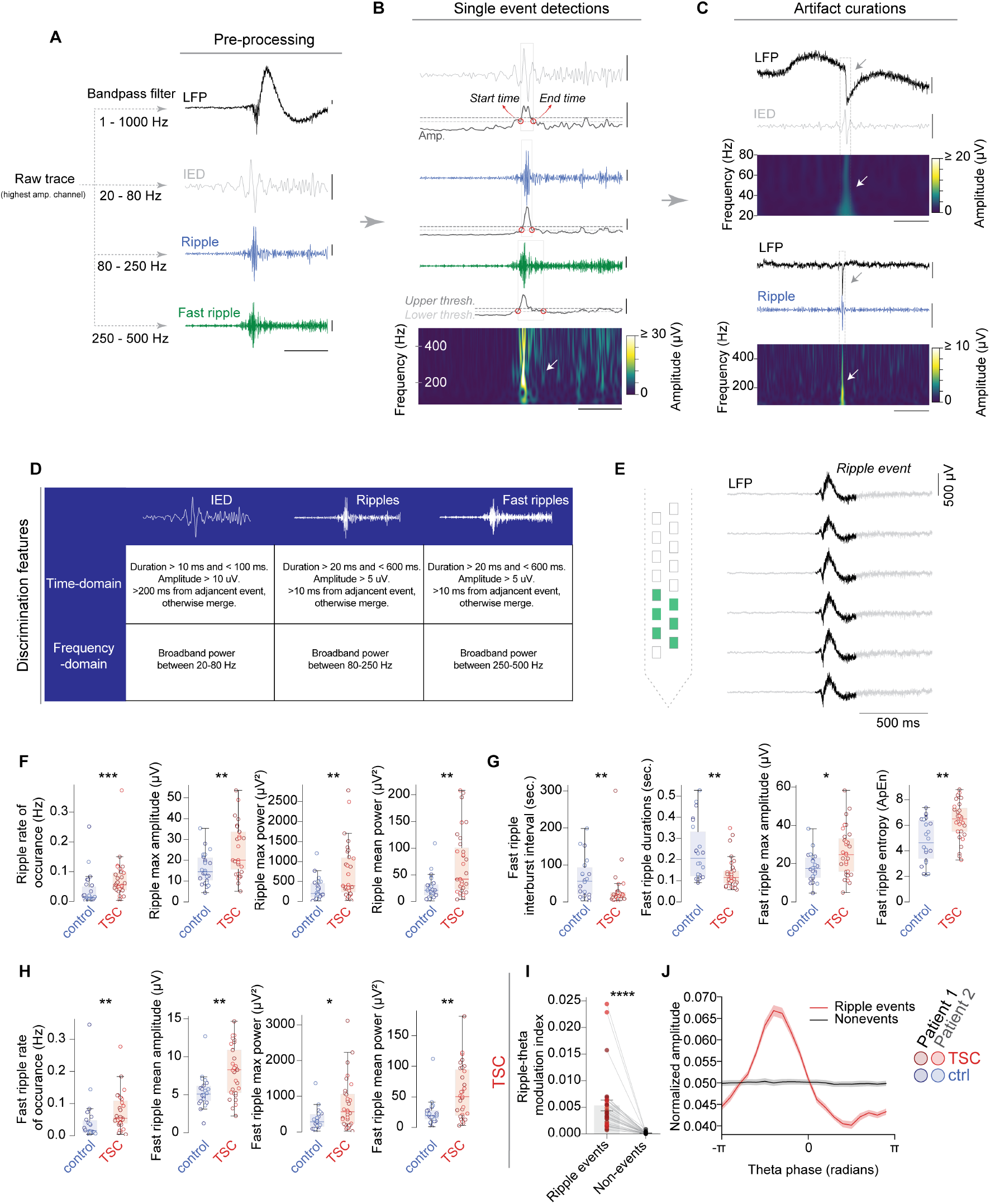
HFOs in TSC have pathological features, related to Figure 3. (A), (B), and (C), Schematic of signal processing pipeline for extracellular recordings in cerebral brain organoids and IED, and high-frequency oscillation detections (see Methods). (B) Example of IED, ripple, and fast ripple event detection with upper and lower detection thresholds marked. (C) Example signal artifacts excluded during the analysis, including a 20-80 Hz (top) and 80-250 Hz example (bottom). Scale bars: 200 ms (time, horizontal), 20 µV (amplitude, vertical). (D) Summary of discrimination features used to classify IEDs, ripples and fast ripples in extracellular recordings from cerebral organoids. (E), In TSC organoids, large field potentials were detected across multiple neighboring electrodes during ripple events. (F), Additional quantifications of ripple signal features in control and TSC brain organoids. In TSC, ripples were shorter and more frequent with increased amplitude and power (control, n = 26 organoids from 11 differentiation experiments; TSC, n = 31 organoids from 11 differentiation experiments; rate of occurrence, P = 0.0006; max amplitude, P = 0.008; max power, P = 0.0057; mean power, P = 0.0016) (G) and (H), Similar to ripples, fast ripples are dysregulated in TSC organoids with characteristically lower inter-burst interval (control, n = 22 organoids from 10 differentiation experiments; TSC, n = 31 organoids from 11 differentiation experiments; P = 0.0036), lower durations (P = 0.0011), higher maximum amplitude (P = 0.016), and higher entropy (C, P = 0.0016; matched to ripples in Fig. 3C to F). Furthermore, fast-ripple rate of occurrence (P = 0.0039), mean amplitude (P = 0.003), maximum power (P = 0.0124), and mean power (P = 0.0032) are increased (D). ApEN, approximate entropy. (I) Distribution of HFO amplitudes in single organoid examples across theta phases reveals strong modulation only within (solid lines), but not outside of ripple events (dashed lines). (J) HFO-theta modulation index analysis shows consistent modulation only in ripple events, compared with periods outside of events (TSC, n = 31 organoids from 11 differentiation experiments, P < 0.0001; Wilcoxon signed-rank test). Within ripple events, TSC organoids show significantly stronger modulation. Boxplots shown in (F), (G) and (H): center, median; lower hinge, 25% quantile; upper hinge, 75% quantile; whiskers ± 1.5 x interquartile range. Data shown in (I) and (J) are presented as mean ± SEM.*P<0.05; ***P<0.001; ****P<0.0001; Mann-Whitney test. For (F) to (I), colors correspond to patient-derived iPSC cell-lines.

**Fig. S4.**
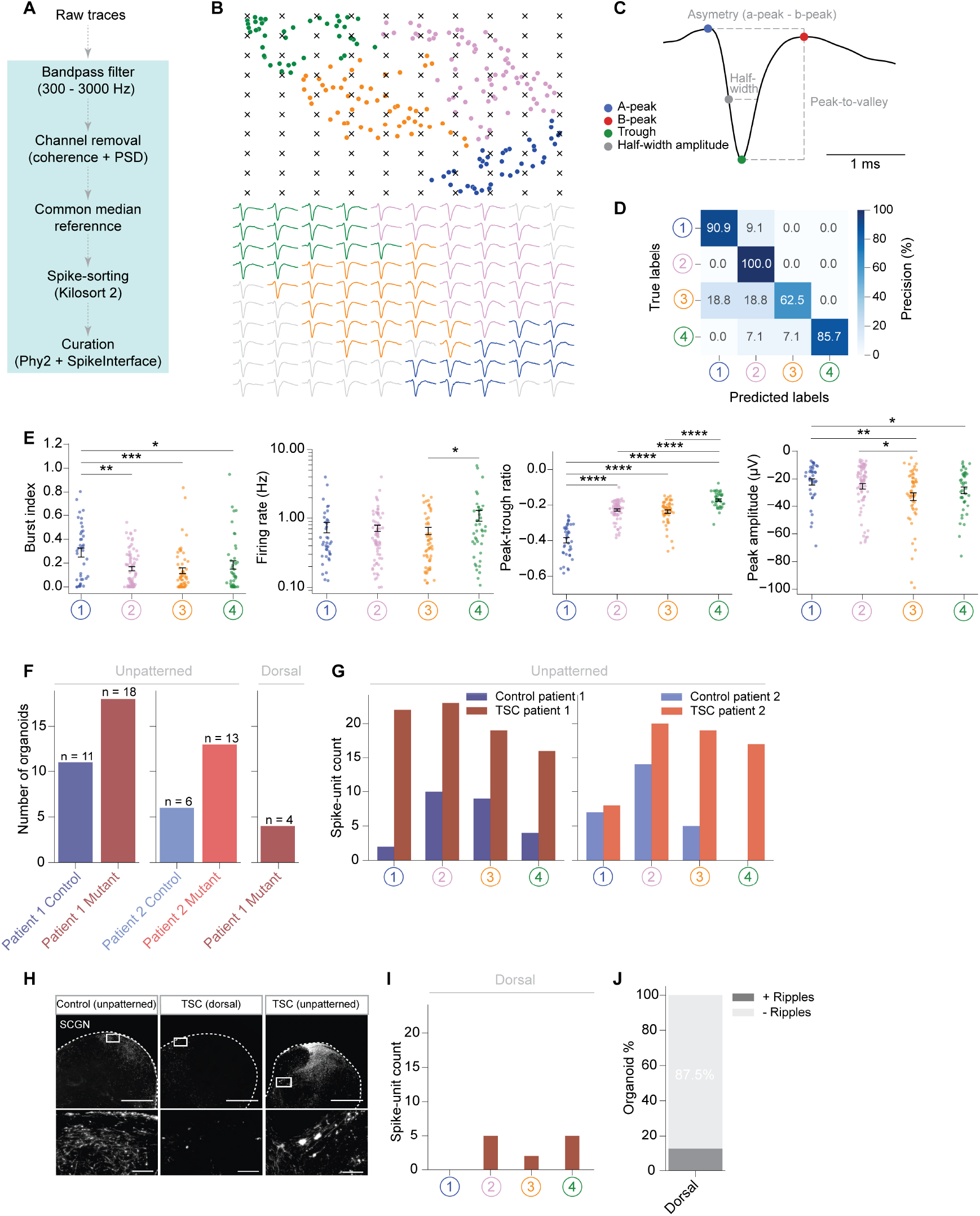
Spike-sorting reveals distinct waveform clusters, related to Figure 4. (A) Processing pipeline for spike sorting in control and TSC organoids using Cambridge Neurotech 64-channel silicon probes. (B) Top: WaveMAP waveform clustering applied to extracellular action potentials (Methods), visualized in a UMAP embedded space (Fig. 3b). A grid of test points (black x’s) is overlaid, spanning the UMAP embedding. Bottom: Inverse UMAP transform applied to the test points, showcasing the predicted waveforms associated with each test point. The visualization highlights the changes in extracellular waveform features across the embedded space. (C) Illustration of waveform features used for analyses. (D) Confusion matrix for a gradient boosted decision tree classifier with 5-fold cross-validation and hyperparamter optimization, performed on waveform clusters 1-4 from 207 cells across organoids from control (n = 17 organoids from 8 differentiation experiments) and TSC (n = 35 organoids from 14 differentiation experiments) experimental groups. The precision of classifications across waveform clusters 1-4 are shown. (E) Quantification of burst-index, firing rate, peak-to-trough ratio and peak amplitude across waveform clusters 1-4. Boxplots shown as the following: center, median; lower hinge, 25% quantile; upper hinge, 75% quantile; whiskers ± 1.5 x interquartile range. Data presented as mean ± SEM.*P<0.05; ***P<0.001; ****P<0.0001; Mann-Whitney test. (F) Total number of control (n = 17 organoids from 8 differentiation experiments) and TSC organoids (n = 35 from 14 differentiation experiments) and (G) corresponding extracted spike-units across unpatterned and dorsal protocols. (H) Immunostainings of the CGE marker SCGN on isogenic control organoids with dorsalized patterning (left) or unpatterned organoids containing both excitatory-inhibitory neurons (middle left), and on TSC organoids with dorsalized patterning (middle right) or unpatterned (right). In both unpatterned organoids there are many more SCGN+ CGE interneurons than in the dorsal counterpart. Scale bar: 1 ms (I) Number of single units extracted for each waveform cluster from TSC organoids (n = 4). (J) Quantification of the percentage of dorsalized organoids that have ripples.

**Table S1.** Clinical characteristics of children with TSC who underwent ioECoG-tailored surgery, related to Figure 2. Demographic information and seizure outcome of included children with TSC.

| Patient | Sex | Age at surgery (y) | Mutation | Surgical side | Longterm Seizure Outcome (4y post-surgery) |
| --- | --- | --- | --- | --- | --- |
| 1 | m | 5.6 | TSC1 (VUS) | Right | ILAE class 1<br>(Note: AED completely withdrawn) |
| 2 | f | 2.2 | TSC1 | Right | ILAE class 1<br>(Note: AED completely withdrawn) |

## References

1 Ethan M. Goldberg and Douglas A. Coulter. Mechanisms of epileptogenesis: a convergence on neural circuit dysfunction. Nature Reviews Neuroscience, 14(5):337–349, 2013. ISSN 1471-003X. doi: 10.1038/nrn3482.

2 Orrin Devinsky, Annamaria Vezzani, Terence J. O’Brien, Nathalie Jette, Ingrid E. Scheffer, Marco de Curtis, and Piero Perucca. Epilepsy. Nature Reviews Disease Primers, 4(1): 18024, 2018. doi: 10.1038/nrdp.2018.24.

3 Wolfgang Löscher. Animal models of seizures and epilepsy: Past, present, and future role for the discovery of antiseizure drugs. Neurochemical Research, 42(7):1873–1888, 2017. ISSN 0364-3190. doi: 10.1007/s11064-017-2222-z.

4 Jim Berg, Staci A. Sorensen, Jonathan T. Ting, Jeremy A. Miller, Thomas Chartrand, Anatoly Buchin, Trygve E. Bakken, Agata Budzillo, Nick Dee, Song-Lin Ding, Nathan W. Gouwens, Rebecca D. Hodge, Brian Kalmbach, Changkyu Lee, Brian R. Lee, Lauren Alfiler, Katherine Baker, Eliza Barkan, Allison Beller, Kyla Berry, Darren Bertagnolli, Kris Bickley, Jasmine Bomben, Thomas Braun, Krissy Brouner, Tamara Casper, Peter Chong, Kirsten Crichton, Rachel Dalley, Rebecca de Frates, Tsega Desta, Samuel Dingman Lee, Florence D’Orazi, Nadezhda Dotson, Tom Egdorf, Rachel Enstrom, Colin Farrell, David Feng, Olivia Fong, Szabina Furdan, Anna A. Galakhova, Clare Gamlin, Amanda Gary, Alexandra Glandon, Jeff Goldy, Melissa Gorham, Natalia A. Goriounova, Sergey Gratiy, Lucas Graybuck, Hong Gu, Kristen Hadley, Nathan Hansen, Tim S. Heistek, Alex M. Henry, Djai B. Heyer, DiJon Hill, Chris Hill, Madie Hupp, Tim Jarsky, Sara Kebede, Lisa Keene, Lisa Kim, Mean-Hwan Kim, Matthew Kroll, Caitlin Latimer, Boaz P. Levi, Katherine E. Link, Matthew Mallory, Rusty Mann, Desiree Marshall, Michelle Maxwell, Medea McGraw, Delissa McMillen, Erica Melief, Eline J. Mertens, Leona Mezei, Norbert Mihut, Stephanie Mok, Gabor Molnar, Alice Mukora, Lindsay Ng, Kiet Ngo, Philip R. Nicovich, Julie Nyhus, Gaspar Olah, Aaron Oldre, Victoria Omstead, Attila Ozsvar, Daniel Park, Hanchuan Peng, Trangthanh Pham, Christina A. Pom, Lydia Potekhina, Ramkumar Rajanbabu, Shea Ransford, David Reid, Christine Rimorin, Augustin Ruiz, David Sandman, Josef Sulc, Susan M. Sunkin, Aaron Szafer, Viktor Szemenyei, Elliot R. Thomsen, Michael Tieu, Amy Torkelson, Jessica Trinh, Herman Tung, Wayne Wakeman, Femke Waleboer, Katelyn Ward, René Wilbers Grace Williams, Zizhen Yao, Jae-Geun Yoon, Costas Anastassiou, Anton Arkhipov, Pal Barzo, Amy Bernard, Charles Cobbs, Philip C. de Witt Hamer, Richard G. Ellenbogen, Luke Esposito, Manuel Ferreira, Ryder P. Gwinn, Michael J. Hawrylycz, Patrick R. Hof, Sander Idema, Allan R. Jones, C. Dirk Keene, Andrew L. Ko, Gabe J. Murphy, Lydia Ng, Jeffrey G. Ojemann, Anoop P. Patel, John W. Phillips, Daniel L. Silbergeld, Kimberly Smith, Bosiljka Tasic, Rafael Yuste, Idan Segev, Christiaan P. J. de Kock, Huibert D. Mansvelder, Gabor Tamas, Hongkui Zeng, Christof Koch, and Ed S. Lein. Human neocortical expansion involves glutamatergic neuron diversification. Nature, 598(7879):151–158, 2021. ISSN 0028-0836. doi: 10.1038/s41586-021-03813-8.

5 T. J. Nowakowski, A. Bhaduri, A. A. Pollen, B. Alvarado, M. A. Mostajo-Radji, E. Di Lullo, M. Haeussler, C. Sandoval-Espinosa, S. J. Liu, D. Velmeshev, J. R. Ounadjela, J. Shuga, X. Wang, D. A. Lim, J. A. West, A. A. Leyrat, W. J. Kent, and A. R. Kriegstein. Spatiotemporal gene expression trajectories reveal developmental hierarchies of the human cortex. Science, 358(6368):1318–1323, 2017. ISSN 1095-9203. doi: 10.1126/science.aap8809.

6 Sahil Loomba, Jakob Straehle, Vijayan Gangadharan, Natalie Heike, Abdelrahman Khalifa, Alessandro Motta, Niansheng Ju, Meike Sievers, Jens Gempt, Hanno S. Meyer, and Moritz Helmstaedter. Connectomic comparison of mouse and human cortex. Science, 2022. ISSN 0036-8075. doi: 10.1126/science.abo0924.

7 Pierre Vanderhaeghen and Franck Polleux. Developmental mechanisms underlying the evolution of human cortical circuits. Nature Reviews Neuroscience, 24(4):213–232, 2023. ISSN 1471-003X. doi: 10.1038/s41583-023-00675-z.

8 GBD 2016 Epilepsy Collaborators, Ettore Beghi, Giorgia Giussani, Emma Nichols, Foad Abd-Allah, Jemal Abdela, Ahmed Abdelalim, Haftom Niguse Abraha, Mina G. Adib, Sutapa Agrawal, Fares Alahdab, Ashish Awasthi, Yohanes Ayele, Miguel A Barboza, Abate Bekele Belachew, Belete Biadgo, Ali Bijani, Helen Bitew, Félix Carvalho, Yazan Chaiah, Ahmad Daryani, Huyen Phuc Do, Manisha Dubey, Aman Yesuf Yesuf Endries, Sharareh Eskandarieh, Andre Faro, Farshad Farzadfar, Seyed-Mohammad Fereshtehnejad, Eduarda Fernandes, Daniel Obadare Fijabi, Irina Filip, Florian Fischer, Abadi Kahsu Gebre, Afewerki Gebremeskel Tsadik, Teklu Gebrehiwo Gebremichael, Kebede Embaye Gezae, Maryam Ghasemi-Kasman, Kidu Gidey Weldegwergs, Meaza Girma Degefa, Elena V. Gnedovskaya, Tekleberhan B Hagos, Arvin Haj-Mirzaian, Arya Haj-Mirzaian, Hamid Yimam Hassen, Simon I Hay, Mihajlo Jakovljevic, Amir Kasaeian, Tesfaye Dessale Kassa, Yousef Saleh Khader, Ibrahim Khalil, Ejaz Ahmad Khan, Jagdish Khubchandani, Adnan Kisa, Kristopher J Krohn, Chanda Kulkarni, Yirga Legesse Nirayo, Mark T Mackay, Marek Majdan, Azeem Majeed, Treh Manhertz, Man Mohan Mehndiratta, Tesfa Mekonen, Hagazi Gebre Meles, Getnet Mengistu, Shafiu Mohammed, Mohsen Naghavi, Ali H Mokdad, Ghulam Mustafa, Seyed Sina Naghibi Irvani, Long Hoang Nguyen, Molly R Nixon, Felix Akpojene Ogbo, Andrew T Olagunju, Tinuke O Olagunju, Mayowa Ojo Owolabi, Michael R Phillips, Gabriel David Pinilla-Monsalve, Mostafa Qorbani, Amir Radfar, Anwar Rafay, Vafa Rahimi-Movaghar, Nickolas Reinig, Perminder S Sachdev, Hosein Safari, Saeed Safari, Saeid Safiri, Mohammad Ali Sahraian, Abdallah M. Samy, Shahabeddin Sarvi, Monika Sawhney, Masood A Shaikh, Mehdi Sharif, Gagandeep Singh, Mari Smith, Cassandra E I Szoeke, Rafael Tabarés-Seisdedos, Mohamad-Hani Temsah, Omar Temsah, Miguel Tortajada-Girbés, Bach Xuan Tran, Amanuel Amanuel Tesfay Tsegay, Irfan Ullah, Narayanaswamy Venketasubramanian, Ronny Westerman, Andrea Sylvia Winkler, Ebrahim M Yimer, Naohiro Yonemoto, Valery L. Feigin, Theo Vos, and Christopher J L Murray. Global, regional, and national burden of epilepsy, 1990–2016: a systematic analysis for the global burden of disease study 2016. The Lancet Neurology, 18(4):357–375, 2019. ISSN 1474-4422. doi:10.1016/s1474-4422(18)30454-x.

9 Patrick Kwan and Martin J. Brodie. Early identification of refractory epilepsy. The New England Journal of Medicine, 342(5):314–319, 2000. ISSN 0028-4793. doi: 10.1056/nejm200002033420503.

10 Zhibin Chen, Martin J. Brodie, Danny Liew, and Patrick Kwan. Treatment outcomes in patients with newly diagnosed epilepsy treated with established and new antiepileptic drugs: A 30-year longitudinal cohort study. JAMA Neurology, 75(3):279, 2017. ISSN 2168-6149. doi: 10.1001/jamaneurol.2017.3949.

11 Oliver L. Eichmüller and Juergen A. Knoblich. Human cerebral organoids — a new tool for clinical neurology research. Nature Reviews Neurology, pages 1–20, 2022. ISSN 1759-4758. doi: 10.1038/s41582-022-00723-9.

12 Kevin W Kelley and Sergiu P Paşca. Human brain organogenesis: Toward a cellular understanding of development and disease. Cell, 185(1):42–61, 2022. ISSN 0092-8674. doi:10.1016/j.cell.2021.10.003.

13 Silvia Velasco, Bruna Paulsen, and Paola Arlotta. 3d brain organoids: Studying brain development and disease outside the embryo. Annual Review of Neuroscience, 43(1):375–389, 2020. ISSN 0147-006X. doi: 10.1146/annurev-neuro-070918-050154.

14 Tal Sharf, Tjitse van der Molen, Stella M. K. Glasauer, Elmer Guzman, Alessio P. Buccino, Gabriel Luna, Zhuowei Cheng, Morgane Audouard, Kamalini G. Ranasinghe, Kiwamu Kudo, Srikantan S. Nagarajan, Kenneth R. Tovar, Linda R. Petzold, Andreas Hierlemann, Paul K. Hansma, and Kenneth S. Kosik. Functional neuronal circuitry and oscillatory dynamics in human brain organoids. Nature Communications, 13(1):4403, 2022. doi:10.1038/s41467-022-32115-4.

15 Cleber A Trujillo, Richard Gao, Priscilla D Negraes, Jing Gu, Justin Buchanan, Sebastian Preissl, Allen Wang, Wei Wu, Gabriel G Haddad, Isaac A Chaim, Alain Domissy, Matthieu Vandenberghe, Anna Devor, Gene W Yeo, Bradley Voytek, and Alysson R Muotri. Complex oscillatory waves emerging from cortical organoids model early human brain network development. Cell stem cell, 25(4):558 – 569.e7, 10 2019. ISSN 1934-5909. doi:10.1016/j.stem.2019.08.002.

16 Ranmal A. Samarasinghe, Osvaldo A. Miranda, Jessie E. Buth, Simon Mitchell, Isabella Ferando, Momoko Watanabe, Thomas F. Allison, Arinnae Kurdian, Namie N. Fotion, Michael J. Gandal, Peyman Golshani, Kathrin Plath, William E. Lowry, Jack M. Parent, Istvan Mody, and Bennett G. Novitch. Identification of neural oscillations and epileptiform changes in human brain organoids. Nature Neuroscience, pages 1–13, 2021. ISSN 1097-6256. doi: 10.1038/s41593-021-00906-5.

17 Hongyi Ye, Cong Chen, Shennan A. Weiss, and Shuang Wang. Pathological and physiological high-frequency oscillations on electroencephalography in patients with epilepsy. Neuroscience Bulletin, 40(5):609–620, 2024. ISSN 1673-7067. doi: 10.1007/s12264-023-01150-6.

18 Oliver L. Eichmüller, Nina S. Corsini, Ábel Vértesy, Ilaria Morassut, Theresa Scholl, Victoria-Elisabeth Gruber, Angela M. Peer, Julia Chu, Maria Novatchkova, Johannes A. Hainfellner, Mercedes F. Paredes, Martha Feucht, and Jürgen A. Knoblich. Amplification of human interneuron progenitors promotes brain tumors and neurological defects. Science, 375(6579), 2022. ISSN 0036-8075. doi: 10.1126/science.abf5546.

19 E. P. Henske, S. Jozwiak, J. C. Kingswood, J. R. Sampson, and E. A. Thiele. Tuberous sclerosis complex. Nat Rev Dis Primers, 2(1):16035, 2016. ISSN 2056-676X. doi: 10.1038/nrdp.2016.35.

20 Catherine J. Chu-Shore, Philippe Major, Susana Camposano, David Muzykewicz, and Elizabeth A. Thiele. The natural history of epilepsy in tuberous sclerosis complex. Epilepsia, 51(7):1236–1241, 2010. ISSN 0013-9580. doi: 10.1111/j.1528-1167.2009.02474.x.

21 A. B. Gelot and A. Represa. Progression of fetal brain lesions in tuberous sclerosis complex. Front Neurosci, 14:899, 2020. ISSN 1662-4548. doi: 10.3389/fnins.2020.00899.

22 Véronique Ruppe, Pelin Dilsiz, Carol Shoshkes Reiss, Chad Carlson, Orrin Devinsky, David Zagzag, Howard L. Weiner, and Delia M. Talos. Developmental brain abnormalities in tuberous sclerosis complex: A comparative tissue analysis of cortical tubers and perituberal cortex. Epilepsia, 55(4):539–550, 2014. ISSN 0013-9580. doi: 10.1111/epi.12545.

23 Lakshminarayanan Kannan, Simon Vogrin, Catherine Bailey, Wirginia Maixner, and A Simon Harvey. Centre of epileptogenic tubers generate and propagate seizures in tuberous sclerosis. Brain, 139(10):2653–2667, 2016. ISSN 0006-8950. doi: 10.1093/brain/aww192.

24 Ahmad R. Mohamed, Catherine A. Bailey, Jeremy L. Freeman, Wirginia Maixner, Graeme D. Jackson, and A. Simon Harvey. Intrinsic epileptogenicity of cortical tubers revealed by intracranial EEG monitoring. Neurology, 79(23):2249–2257, 2012. ISSN 0028-3878. doi: 10.1212/wnl.0b013e3182768923.

25 Elodie Despouy, Jonathan Curot, Marie Denuelle, Martin Deudon, Jean-Christophe Sol, Jean-Albert Lotterie, Leila Reddy, Lionel G. Nowak, Jérémie Pariente, Simon J. Thorpe, Luc Valton, and Emmanuel J. Barbeau. Neuronal spiking activity highlights a gradient of epileptogenicity in human tuberous sclerosis lesions. Clinical Neurophysiology, 130(4):537–547, 2019. ISSN 1388-2457. doi: 10.1016/j.clinph.2018.12.013.

26 Nicola Specchio, Giusy Carfi Pavia, Luca de Palma, Alessandro De Benedictis, Chiara Pepi, Marta Conti, Carlo Efisio Marras, Federico Vigevano, and Paolo Curatolo. Current role of surgery for tuberous sclerosis complex-associated epilepsy. Pediatric Investigation, 6(1): 16–22, 2022. ISSN 2096-3726. doi: 10.1002/ped4.12312.

27 Richard J. Staba, Charles L. Wilson, Anatol Bragin, Donald Jhung, Itzhak Fried, and Jerome Engel. High-frequency oscillations recorded in human medial temporal lobe during sleep. Annals of Neurology, 56(1):108–115, 2004. ISSN 0364-5134. doi: 10.1002/ana.20164.

28 Maeike Zijlmans, Willemiek Zweiphenning, and Nicole van Klink. Changing concepts in presurgical assessment for epilepsy surgery. Nature Reviews Neurology, 15(10):594–606, 2019. ISSN 1759-4758. doi: 10.1038/s41582-019-0224-y.

29 Willemiek Zweiphenning, Maryse A van ‘t Klooster, Nicole E C van Klink, Frans S S Leijten, Cyrille H Ferrier, Tineke Gebbink, Geertjan Huiskamp, Martine JE van Zandvoort, Monique M J van Schooneveld, M Bourez, Sophie Goemans, Sven Straumann, Peter C van Rijen, Peter H Gosselaar, Pieter van Eijsden, Willem M Otte, Eric van Diessen, Kees PJ Braun, Maeike Zijlmans, HFO study group, Eltje M. Bloemen-Carlier, Veronika Cibulková, Renee de Munnink, Sandra van der Salm, Martinus J.C. Eijkemans, Janine M. Ophorst-van Eck, Anouk Velders, Charlotte J.J. van Asch, Jack Zwemmer, Renate van Regteren-van Griethuysen, Henriette Smeding, Lydia van der Berg, Jeroen de Bresser, Gérard A.P. de Kort, and Jan-Willem Dankbaar. Intraoperative electrocorticography using high-frequency oscillations or spikes to tailor epilepsy surgery in the netherlands (the HFO trial): a randomised, single-blind, adaptive non-inferiority trial. The Lancet Neurology, 21(11):982–993, 2022. ISSN 1474-4422. doi: 10.1016/s1474-4422(22)00311-8.

30 Floor E. Jansen, Alexander. C. Van Huffelen, ’Ále Algra, and Onno Van Nieuwenhuizen. Epilepsy surgery in tuberous sclerosis: A systematic review. Epilepsia, 48(8):1477–1484, 2007. ISSN 0013-9580. doi: 10.1111/j.1528-1167.2007.01117.x.

31 M Zijlmans, J Jacobs, R Zelmann, F Dubeau, and J Gotman. High-frequency oscillations mirror disease activity in patients with epilepsy. Neurology, 72(11):979–986, 2009. ISSN 0028-3878. doi: 10.1212/01.wnl.0000344402.20334.81.

32 Xiaolong Zou and Da-Hui Wang. On the phase relationship between excitatory and inhibitory neurons in oscillation. Frontiers in Computational Neuroscience, 10:138, 2016. ISSN 1662-5188. doi: 10.3389/fncom.2016.00138.

33 Cedric Bardy, Mark van den Hurk, Tameji Eames, Cynthia Marchand, Ruben V. Hernandez, Mariko Kellogg, Mark Gorris, Ben Galet, Vanessa Palomares, Joshua Brown, Anne G. Bang, Jerome Mertens, Lena Böhnke, Leah Boyer, Suzanne Simon, and Fred H. Gage. Neuronal medium that supports basic synaptic functions and activity of human neurons in vitro. Proc Natl Acad Sci U S A, 112(25):E3312, 2015. ISSN 1091-6490. doi: 10.1073/pnas.1509741112.

34 Y. Ben-Ari, I. Khalilov, K. T. Kahle, and E. Cherubini. The GABA excitatory/inhibitory shift in brain maturation and neurological disorders. Neuroscientist, 18(5):467–86, 2012. ISSN 1089-4098. doi: 10.1177/1073858412438697.

35 Richard Gao, Michael Deistler, Auguste Schulz, Pedro J. Gonçalves, and Jakob H. Macke. Deep inverse modeling reveals dynamic-dependent invariances in neural circuit mechanisms. bioRxiv, page 2024.08.21.608969, 2024. doi: 10.1101/2024.08.21.608969.

36 Bassam V. Atallah and Massimo Scanziani. Instantaneous modulation of gamma oscillation frequency by balancing excitation with inhibition. Neuron, 62(4):566–577, 2009. ISSN 0896-6273. doi: 10.1016/j.neuron.2009.04.027.

37 Richard Gao, Erik J. Peterson, and Bradley Voytek. Inferring synaptic excitation/inhibition balance from field potentials. NeuroImage, 158:70–78, 2017. ISSN 1053-8119. doi: 10.1016/j.neuroimage.2017.06.078.

38 Anthony A. Oliva, Trang T. Lam, and John W. Swann. Distally directed dendrotoxicity induced by kainic acid in hippocampal interneurons of green fluorescent protein-expressing transgenic mice. The Journal of Neuroscience, 22(18):8052–8062, 2002. ISSN 0270-6474. doi: 10.1523/jneurosci.22-18-08052.2002.

39 M. Wong and D. Guo. Dendritic spine pathology in epilepsy: Cause or consequence? Neuroscience, 251:141–150, 2013. ISSN 0306-4522. doi: 10.1016/j.neuroscience.2012.03.048.

40 Peter R. Huttenlocher and Peter T. Heydemann. Fine structure of cortical tubers in tuberous sclerosis: A golgi study. Annals of Neurology, 16(5):595–602, 1984. ISSN 0364-5134. doi: 10.1002/ana.410160511.

41 Laura Rossini, Dalia De Santis, Roberta Rosa Mauceri, Chiara Tesoriero, Marina Bentivoglio, Emanuela Maderna, Antonio Maiorana, Francesco Deleo, Marco de Curtis, Giovanni Tringali, Massimo Cossu, Gemma Tumminelli, Manuela Bramerio, Roberto Spreafico, Laura Tassi, and Rita Garbelli. Dendritic pathology, spine loss and synaptic reorganization in human cortex from epilepsy patients. Brain, 144(1):251–265, 2020. ISSN 0006-8950. doi: 10.1093/brain/awaa387.

42 Maxime Lévesque, Pariya Salami, Zahra Shiri, and Massimo Avoli. Interictal oscillations and focal epileptic disorders. European Journal of Neuroscience, 48(8):2915–2927, 2018. ISSN 0953-816X. doi: 10.1111/ejn.13628.

43 Julia Jacobs, Joyce Y Wu, Piero Perucca, Rina Zelmann, Malenka Mader, Francois Dubeau, Gary W Mathern, Andreas Schulze-Bonhage, and Jean Gotman. Removing high-frequency oscillations: A prospective multicenter study on seizure outcome. Neurology, 91(11):e1040– e1052, 2018. ISSN 0028-3878. doi: 10.1212/wnl.0000000000006158.

44 Ziyi Wang, Jiaojiao Guo, Maryse van ‘t Klooster, Sem Hoogteijling, Julia Jacobs, and Maeike Zijlmans. Prognostic value of complete resection of the high-frequency oscillation area in intracranial EEG. Neurology, 102(9):e209216, 2024. ISSN 0028-3878. doi: 10.1212/wnl.0000000000209216.

45 Alex P Vaz, Sara K Inati, Nicolas Brunel, and Kareem A Zaghloul. Coupled ripple oscillations between the medial temporal lobe and neocortex retrieve human memory. Science (New York, N.Y.), 363(6430):975–978, 2019. ISSN 0036-8075. doi: 10.1126/science.aau8956.

46 Sergey Burnos, Birgit Frauscher, Rina Zelmann, Claire Haegelen, Johannes Sarnthein, and Jean Gotman. The morphology of high frequency oscillations (HFO) does not improve delineating the epileptogenic zone. Clinical Neurophysiology, 127(4):2140–2148, 2016. ISSN 1388-2457. doi: 10.1016/j.clinph.2016.01.002.

47 Ai Phuong S Tong, Alex P Vaz, John H Wittig, Sara K Inati, and Kareem A Zaghloul. Ripples reflect a spectrum of synchronous spiking activity in human anterior temporal lobe. eLife, 10:e68401, 2021. doi: 10.7554/elife.68401.

48 Alessandra Maccabeo, Maryse A. van ‘t Klooster, Eline Schaft, Matteo Demuru, Willemiek Zweiphenning, Peter Gosselaar, Tineke Gebbink, Wim M. Otte, and Maeike Zijlmans. Spikes and high frequency oscillations in lateral neocortical temporal lobe epilepsy: Can they predict the success chance of hippocampus-sparing resections? Frontiers in Neurology, 13:797075, 2022. ISSN 1664-2295. doi: 10.3389/fneur.2022.797075.

49 Shuang Wang, Irene Z. Wang, Juan C. Bulacio, John C. Mosher, Jorge Gonzalez-Martinez, Andreas V. Alexopoulos, Imad M. Najm, and Norman K. So. Ripple classification helps to localize the seizure-onset zone in neocortical epilepsy. Epilepsia, 54(2):370–376, 2013. ISSN 0013-9580. doi: 10.1111/j.1528-1167.2012.03721.x.

50 Birgit Frauscher, Nicolás von Ellenrieder, Rina Zelmann, Christine Rogers, Dang Khoa Nguyen, Philippe Kahane, François Dubeau, and Jean Gotman. High-frequency oscillations in the normal human brain. Annals of Neurology, 84(3):374–385, 2018. ISSN 0364-5134. doi: 10.1002/ana.25304.

51 Marisol Soula, Anna Maslarova, Ryan E. Harvey, Manuel Valero, Sebastian Brandner, Hajo Hamer, Antonio Fernández-Ruiz, and György Buzsáki. Interictal epileptiform discharges affect memory in an alzheimer’s disease mouse model. Proceedings of the National Academy of Sciences, 120(34):e2302676120, 2023. ISSN 0027-8424. doi: 10.1073/pnas.2302676120.

52 Yutaka Nonoda, Makoto Miyakoshi, Alejandro Ojeda, Scott Makeig, Csaba Juhász, Sandeep Sood, and Eishi Asano. Interictal high-frequency oscillations generated by seizure onset and eloquent areas may be differentially coupled with different slow waves. Clinical Neurophysiology, 127(6):2489–2499, 2016. ISSN 1388-2457. doi: 10.1016/j.clinph.2016.03.022.

53 Mina Amiri, Birgit Frauscher, and Jean Gotman. Phase-amplitude coupling is elevated in deep sleep and in the onset zone of focal epileptic seizures. Frontiers in Human Neuroscience, 10:387, 2016. ISSN 1662-5161. doi: 10.3389/fnhum.2016.00387.

54 Hiroaki Hashimoto, Hui Ming Khoo, Takufumi Yanagisawa, Naoki Tani, Satoru Oshino, Haruhiko Kishima, and Masayuki Hirata. Phase-amplitude coupling of ripple activities during seizure evolution with theta phase. Clinical Neurophysiology, 132(6):1243–1253, 2021. ISSN 1388-2457. doi: 10.1016/j.clinph.2021.03.007.

55 Inkyung Song, Iren Orosz, Inna Chervoneva, Zachary J. Waldman, Itzhak Fried, Chengyuan Wu, Ashwini Sharan, Noriko Salamon, Richard Gorniak, Sandra Dewar, Anatol Bragin, Jerome Engel, Michael R. Sperling, Richard Staba, and Shennan A. Weiss. Bimodal coupling of ripples and slower oscillations during sleep in patients with focal epilepsy. Epilepsia, 58(11):1972–1984, 2017. ISSN 0013-9580. doi: 10.1111/epi.13912.

56 Alessio P Buccino, Cole L Hurwitz, Samuel Garcia, Jeremy Magland, Joshua H Siegle, Roger Hurwitz, and Matthias H Hennig. SpikeInterface, a unified framework for spike sorting. eLife, 9:e61834, 2020. doi: 10.7554/elife.61834.

57 György Buzsáki, Costas A. Anastassiou, and Christof Koch. The origin of extracellular fields and currents — EEG, ECoG, LFP and spikes. Nature Reviews Neuroscience, 13(6):407–420, 2012. ISSN 1471-003X. doi: 10.1038/nrn3241.

58 Eric Kenji Lee, Hymavathy Balasubramanian, Alexandra Tsolias, Stephanie Udochukwu Anakwe, Maria Medalla, Krishna V Shenoy, and Chandramouli Chandrasekaran. Non-linear dimensionality reduction on extracellular waveforms reveals cell type diversity in premotor cortex. eLife, 10:e67490, 2021. doi: 10.7554/elife.67490.

59 Kenji Lee, Nicole Carr, Alec Perliss, and Chandramouli Chandrasekaran. WaveMAP for identifying putative cell types from in vivo electrophysiology. STAR Protocols, 4(2):102320, 2023. ISSN 2666-1667. doi: 10.1016/j.xpro.2023.102320.

60 Katharina T. Hofer, Ágnes Kandrács, Kinga Tóth, Boglárka Hajnal, Virág Bokodi, Estilla Zsófia Tóth, Loránd Erő ss László Entz Attila G. Bagó, Dániel Fabó, István Ulbert, and Lucia Wittner. Bursting of excitatory cells is linked to interictal epileptic discharge generation in humans. Scientific Reports, 12(1):6280, 2022. doi: 10.1038/s41598-022-10319-4.

61 Jozsef Csicsvari, Hajime Hirase, András Czurkó, Akira Mamiya, and György Buzsáki. Oscillatory coupling of hippocampal pyramidal cells and interneurons in the behaving rat. The Journal of Neuroscience, 19(1):274–287, 1999. ISSN 0270-6474. doi: 10.1523/jneurosci.19-01-00274.1999.

62 Jozsef Csicsvari, Hajime Hirase, Andras Czurko, and György Buzsáki. Reliability and state dependence of pyramidal cell–interneuron synapses in the hippocampus an ensemble approach in the behaving rat. Neuron, 21(1):179–189, 1998. ISSN 0896-6273. doi: 10.1016/s0896-6273(00)80525-5.

63 Brian R. Lee, Rachel Dalley, Jeremy A. Miller, Thomas Chartrand, Jennie Close, Rusty Mann, Alice Mukora, Lindsay Ng, Lauren Alfiler, Katherine Baker, Darren Bertagnolli, Krissy Brouner, Tamara Casper, Eva Csajbok, Nicholas Donadio, Stan L.W. Driessens, Tom Egdorf, Rachel Enstrom, Anna A. Galakhova, Amanda Gary, Emily Gelfand, Jeff Goldy, Kristen Hadley, Tim S. Heistek, Dijon Hill, Wen-Hsien Hou, Nelson Johansen, Nik Jorstad, Lisa Kim, Agnes Katalin Kocsis, Lauren Kruse, Michael Kunst, Gabriela León, Brian Long, Matthew Mallory, Michelle Maxwell, Medea McGraw, Delissa McMillen, Erica J. Melief, Gabor Molnar, Marty T. Mortrud, Dakota Newman, Julie Nyhus, Ximena Opitz-Araya, Attila Ozsvár, Trangthanh Pham, Alice Pom, Lydia Potekhina, Ram Rajanbabu, Augustin Ruiz, Susan M. Sunkin, Ildikó Szöts, Naz Taskin, Bargavi Thyagarajan, Michael Tieu, Jessica Trinh, Sara Vargas, David Vumbaco, Femke Waleboer, Sarah Walling-Bell, Natalie Weed, Grace Williams, Julia Wilson, Shenqin Yao, Thomas Zhou, Pál Barzó, Trygve Bakken, Charles Cobbs, Nick Dee, Richard G. Ellenbogen, Luke Esposito, Manuel Ferreira, Nathan W. Gouwens, Benjamin Grannan, Ryder P. Gwinn, Jason S. Hauptman, Rebecca Hodge, Tim Jarsky, C. Dirk Keene, Andrew L. Ko, Anders Rosendal Korshoej, Boaz P. Levi, Kaare Meier, Jeffrey G. Ojemann, Anoop Patel, Jacob Ruzevick, Daniel L. Silbergeld, Kimberly Smith, Jens Christian Sørensen, Jack Waters, Hongkui Zeng, Jim Berg, Marco Capogna, Natalia A. Goriounova, Brian Kalmbach, Christiaan P.J. de Kock, Huib D. Mansvelder, Staci A. Sorensen, Gabor Tamas, Ed S. Lein, and Jonathan T. Ting. Signature morphoelectric properties of diverse GABAergic interneurons in the human neocortex. Science, 382(6667): eadf6484, 2023. ISSN 0036-8075. doi: 10.1126/science.adf6484.

64 C Bardy, M van den Hurk, B Kakaradov, JA Erwin, BN Jaeger, RV Hernandez, T Eames, AA Paucar, M Gorris, C Marchand, R Jappelli, J Barron, AK Bryant, M Kellogg, RS Lasken, Wong, Zabolocki, Eichmüller et al. | Cerebral Organoids Uncover Mechanisms of Neural Activity Changes in Epileptogenesis BPF Rutten, HWM Steinbusch, GW Yeo, and FH Gage. Predicting the functional states of human iPSC-derived neurons with single-cell RNA-seq and electrophysiology. Molecular Psychiatry, 21(11):1573–1588, 2016. ISSN 1359-4184. doi: 10.1038/mp.2016.158.

65 Catherine A. Schevon, Sau K. Ng, Joshua Cappell, Robert R. Goodman, Guy McKhann, Allen Waziri, Almut Branner, Alexandre Sosunov, Charles E. Schroeder, and Ronald G. Emerson. Microphysiology of epileptiform activity in human neocortex. Journal of Clinical Neurophysiology, 25(6):321–330, 2008. ISSN 0736-0258. doi: 10.1097/wnp.0b013e31818e8010.

66 Jonathan Curot, Emmanuel Barbeau, Elodie Despouy, Marie Denuelle, Jean Christophe Sol, Jean Albert Lotterie, Luc Valton, and Adrien Peyrache. Local neuronal excitation and global inhibition during epileptic fast ripples in humans. Brain, 2022. ISSN 0006-8950. doi: 10.1093/brain/awac319.

67 Jerome Engel Jr, Anatol Bragin, Richard Staba, and Istvan Mody. High-frequency oscillations: What is normal and what is not? Epilepsia, 50(4):598–604, 2009. ISSN 0013-9580. doi: 10.1111/j.1528-1167.2008.01917.x.

68 Shennan A. Weiss, Iren Orosz, Noriko Salamon, Stephanie Moy, Linqing Wei, Maryse Van’t Klooster, Robert T. Knight, Ronald M. Harper, Anatol Bragin, Itzhak Fried, Jerome Engel, and Richard J. Staba. Ripples on spikes show increased phase-amplitude coupling in mesial temporal lobe epilepsy seizure-onset zones. Epilepsia, 57(11):1916–1930, 2016. ISSN 0013-9580. doi: 10.1111/epi.13572.

69 György Buzsáki and Fernando Lopes da Silva. High frequency oscillations in the intact brain. Progress in Neurobiology, 98(3):241–249, 2012. ISSN 0301-0082. doi: 10.1016/j.pneurobio.2012.02.004.

70 Anne H. Mooij, Geertjan J.M. Huiskamp, Emmeke Aarts, Cyrille H. Ferrier, Kees P.J. Braun, and Maeike Zijlmans. Accurate differentiation between physiological and pathological ripples recorded with scalp-EEG. Clinical Neurophysiology, 143:172–181, 2022. ISSN 1388-2457. doi: 10.1016/j.clinph.2022.08.014.

71 Premysl Jiruska, Marco de Curtis, John G. R. Jefferys, Catherine A. Schevon, Steven J. Schiff, and Kaspar Schindler. Synchronization and desynchronization in epilepsy: controversies and hypotheses. The Journal of Physiology, 591(4):787–797, 2013. ISSN 0022-3751. doi: 10.1113/jphysiol.2012.239590.

72 Sophie Demont-Guignard, Pascal Benquet, Urs Gerber, Arnaud Biraben, Benoit Martin, and Fabrice Wendling. Distinct hyperexcitability mechanisms underlie fast ripples and epileptic spikes. Annals of Neurology, 71(3):342–352, 2012. ISSN 0364-5134. doi: 10.1002/ana.22610.

73 Anatol Bragin, Istvan Mody, Charles L. Wilson, and Jerome Engel. Local generation of fast ripples in epileptic brain. The Journal of Neuroscience, 22(5):2012–2021, 2002. ISSN 0270-6474. doi: 10.1523/jneurosci.22-05-02012.2002.

74 Gilles Huberfeld, Lucia Wittner, Stéphane Clemenceau, Michel Baulac, Kai Kaila, Richard Miles, and Claudio Rivera. Perturbed chloride homeostasis and GABAergic signaling in human temporal lobe epilepsy. The Journal of Neuroscience, 27(37):9866–9873, 2007. ISSN 0270-6474. doi: 10.1523/jneurosci.2761-07.2007.

75 Thoralf Opitz, Ana D. De Lima, and Thomas Voigt. Spontaneous development of synchronous oscillatory activity during maturation of cortical networks in vitro. Journal of Neurophysiology, 88(5):2196–2206, 2002. ISSN 0022-3077. doi: 10.1152/jn.00316.2002.

76 Fikri Birey, Min-Yin Li, Aaron Gordon, Mayuri V. Thete, Alfredo M. Valencia, Omer Revah, Anca M. Pasça, Daniel H. Geschwind, and Sergiu P. Pasça. Dissecting the molecular basis of human interneuron migration in forebrain assembloids from timothy syndrome. Cell Stem Cell, 2022. ISSN 1934-5909. doi: 10.1016/j.stem.2021.11.011.

77 Maria-Patapia Zafeiriou, Guobin Bao, James Hudson, Rashi Halder, Alica Blenkle, MarieKristin Schreiber, Andre Fischer, Detlev Schild, and Wolfram-Hubertus Zimmermann. Developmental GABA polarity switch and neuronal plasticity in bioengineered neuronal organoids. Nature Communications, 11(1):3791, 2020. doi: 10.1038/s41467-020-17521-w.

78 Hua Hu, Jian Gan, and Peter Jonas. Interneurons. fast-spiking, parvalbumin GABAergic interneurons: from cellular design to microcircuit function. Science (New York, N.Y.), 345 (6196):1255263, 2014. ISSN 0036-8075. doi: 10.1126/science.1255263.

79 X Jiang, M Lachance, and E Rossignol. Involvement of cortical fast-spiking parvalbuminpositive basket cells in epilepsy. Progress in brain research, 226:81–126, 2016. ISSN 0079-6123. doi: 10.1016/bs.pbr.2016.04.012.

80 Beulah Leitch. Parvalbumin interneuron dysfunction in neurological disorders: Focus on epilepsy and alzheimer’s disease. International Journal of Molecular Sciences, 25(10):5549, 2024. doi: 10.3390/ijms25105549.

81 Fenna M Krienen, Melissa Goldman, Qiangge Zhang, Ricardo C H del Rosario, Marta Florio, Robert Machold, Arpiar Saunders, Kirsten Levandowski, Heather Zaniewski, Benjamin Schuman, Carolyn Wu, Alyssa Lutservitz, Christopher D Mullally, Nora Reed, Elizabeth Bien, Laura Bortolin, Marian Fernandez-Otero, Jessica D Lin, Alec Wysoker, James Nemesh, David Kulp, Monika Burns, Victor Tkachev, Richard Smith, Christopher A Walsh, Jordane Dimidschstein, Bernardo Rudy, Leslie S Kean, Sabina Berretta, Gord Fishell, Guoping Feng, and Steven A McCarroll. Innovations present in the primate interneuron repertoire. Nature, 586(7828):262–269, 2020. ISSN 0028-0836. doi: 10.1038/s41586-020-2781-z.

82 Ana Hladnik, Domagoj Džaja, Sanja Darmopil, Nataša Jovanov-Milošević, and Zdravko Petanjek. Spatio-temporal extension in site of origin for cortical calretinin neurons in primates. Frontiers in Neuroanatomy, 8:50, 2014. ISSN 1662-5129. doi: 10.3389/fnana.2014.00050.

83 D. V. Hansen, J. H. Lui, P. Flandin, K. Yoshikawa, J. L. Rubenstein, A. Alvarez-Buylla, and R. Kriegstein. Non-epithelial stem cells and cortical interneuron production in the human ganglionic eminences. Nat Neurosci, 16(11):1576–87, 2013. ISSN 1546-1726. doi: 10.1038/nn.3541.

84 Elena Dossi and Gilles Huberfeld. GABAergic circuits drive focal seizures. Neurobiology of Disease, 180:106102, 2023. ISSN 0969-9961. doi: 10.1016/j.nbd.2023.106102.

85 Behnam Molaee-Ardekani, Pascal Benquet, Fabrice Bartolomei, and Fabrice Wendling. Computational modeling of high-frequency oscillations at the onset of neocortical partial seizures: From ‘altered structure’ to ‘dysfunction’. NeuroImage, 52(3):1109–1122, 2010. ISSN 1053-8119. doi: 10.1016/j.neuroimage.2009.12.049.

86 Premysl Jiruska, Catalina Alvarado-Rojas, Catherine A. Schevon, Richard Staba, William Stacey, Fabrice Wendling, and Massimo Avoli. Update on the mechanisms and roles of high-frequency oscillations in seizures and epileptic disorders. Epilepsia, 58(8):1330–1339, 2017. ISSN 0013-9580. doi: 10.1111/epi.13830.

87 Véronique M. André, Nanping Wu, Irene Yamazaki, Snow T. Nguyen, Robin S. Fisher, Harry V. Vinters, Gary W. Mathern, Michael S. Levine, and Carlos Cepeda. Cytomegalic interneurons: A new abnormal cell type in severe pediatric cortical dysplasia. Journal of Neuropathology & Experimental Neurology, 66(6):491–504, 2007. ISSN 0022-3069. doi:10.1097/01.jnen.0000240473.50661.d8.

88 Jessie De Ridder, Birgit Verhelle, Jan Vervisch, Katrien Lemmens, Katarzyna Kotulska, Romina Moavero, Paolo Curatolo, Bernhard Weschke, Kate Riney, Martha Feucht, Pavel Krsek, Rima Nabbout, Anna C Jansen, Konrad Wojdan, Dorota Domanska-Pakieła, Magdalena Kaczorowska-Frontczak, Christoph Hertzberg, Cyrille H Ferrier, Sharon Samueli, Barbora Benova, Eleonora Aronica, David J Kwiatkowski, Floor E Jansen, Sergiusz Jóźwiak, Lieven Lagae, J Anink, A Benvenuto, M Blazejczyk, A Bongaarts, J Borkowska, D Breuillard, D Chmielewski, M Dabrowska, L Emberti Gialloreti, K Giannikou, J Głowacka-Walas, L Hamieh, A Haręza, H Hulshof, A Iyer, B Janssen, J Jaworski, K Lehmann, A Leusman, N Maćkowiak, J D Mills, A Muehlebner, K Sadowski, C Scheldeman, T Scholl, M Schooneveld, A Sciuto, K Sijko, M Slowinska, A Tempes, M Urbańska, and J van Scheppingen. Early epileptiform EEG activity in infants with tuberous sclerosis complex predicts epilepsy and neurodevelopmental outcomes. Epilepsia, 62(5):1208–1219, 2021. ISSN 0013-9580. doi: 10.1111/epi.16892.

89 Sergiusz Jóźwiak, Katarzyna Kotulska, Dorota Domańska-Pakieła, Barbara Łojszczyk, Małgorzata Syczewska, Dariusz Chmielewski, Dorota Dunin-Wąsowicz, Tomasz Kmieć, Joanna Szymkiewicz-Dangel, Maria Kornacka, Wanda Kawalec, Dariusz Kuczyński, Julita Borkowska, Katarzyna Tomaszek, Elžbieta Jurkiewicz, and Maria Respondek-Liberska. Antiepileptic treatment before the onset of seizures reduces epilepsy severity and risk of mental retardation in infants with tuberous sclerosis complex. European Journal of Paediatric Neurology, 15(5):424–431, 2011. ISSN 1090-3798. doi: 10.1016/j.ejpn.2011.03.010.

90 Joyce Y. Wu, Monisha Goyal, Jurriaan M. Peters, Darcy Krueger, Mustafa Sahin, Hope Northrup, Kit S. Au, Sarah O’Kelley, Marian Williams, Deborah A. Pearson, Ellen Hanson, Anna W. Byars, Jessica Krefting, Mark Beasley, Gary Cutter, Nita Limdi, and E. Martina Bebin. Scalp EEG spikes predict impending epilepsy in TSC infants: A longitudinal observational study. Epilepsia, 60(12):2428–2436, 2019. ISSN 0013-9580. doi: 10.1111/epi.16379.

91 Anne-Elise C. de Groen, Jeffrey Bolton, Ann Marie Bergin, Mustafa Sahin, and Jurriaan M. Peters. The evolution of subclinical seizures in children with tuberous sclerosis complex. Journal of Child Neurology, 34(12):770–777, 2019. ISSN 0883-0738. doi:10.1177/0883073819860640.

92 Katarzyna Kotulska, David J. Kwiatkowski, Paolo Curatolo, Bernhard Weschke, Kate Riney, Floor Jansen, Martha Feucht, Pavel Krsek, Rima Nabbout, Anna C. Jansen, Konrad Wojdan, Kamil Sijko, Jagoda Głowacka-Walas, Julita Borkowska, Krzysztof Sadowski, Dorota Domańska-Pakieła, Romina Moavero, Christoph Hertzberg, Hanna Hulshof, Theresa Scholl, Barbora Benova, Eleonora Aronica, Jessie de Ridder, Lieven Lagae, Sergiusz Jóźwiak, J. Anink, E. Aronica, B. Benova, A. Benvenuto, M. Blazejczyk, A. Bongaerts, J. Borkowska, D. Breuillard, D. Chmielewski, P. Curatolo, M. Dabrowska, D. Domańska-Pakieła, M. Feucht, K. Giannikou, J. Głowacka-Walas, L. Hamieh, A. Harza, Ch. Hertzberg, H. Hulshof, F. Huschner, A. Iyer, A. Jansen, F. Jansen, B. Janssen, J. Jaworski, S. Jùźwiak, M. Kaczorowska-Frontczak, K. Kotulska, P. Krsek, D. Kwiatkowski, L. Lagae, K. Lehmann, A. Leusman, N. Maćkowiak, J. Mills, R. Moavero, A. Muelebner, R. Nabbout, J. de Ridder, K. Riney, K. Sadowski, S. Samueli, C. Scheldeman, T. Scholl, A. Sciuto, K. Sijko, M. Słowińska, A. Tempes, J. van Scheppingen, B. Verhelle, J. Vervisch, M. Urbańska, B. Weschke, and K. Wojdan. Prevention of epilepsy in infants with tuberous sclerosis complex in the EPISTOP trial. Annals of Neurology, 89(2):304–314, 2021. ISSN 0364-5134. doi: 10.1002/ana.25956.

93 Nicola Specchio, Rima Nabbout, Eleonora Aronica, Stephane Auvin, Arianna Benvenuto, Luca de Palma, Martha Feucht, Floor Jansen, Katarzyna Kotulska, Harvey Sarnat, Lieven Lagae, Sergiusz Jozwiak, and Paolo Curatolo. Updated clinical recommendations for the management of tuberous sclerosis complex associated epilepsy. European Journal of Paediatric Neurology, 47:25–34, 2023. ISSN 1090-3798. doi: 10.1016/j.ejpn.2023.08.005.

94 E. Martina Bebin, Jurriaan M. Peters, Brenda E. Porter, Tarrant O. McPherson, Sarah O’Kelley, Mustafa Sahin, Katherine S. Taub, Rajsekar Rajaraman, Stephanie C. Randle, William M. McClintock, Mary Kay Koenig, Mike D. Frost, Hope A. Northrup, Klaus Werner, Danielle A. Nolan, Michael Wong, Jessica L. Krefting, Fred Biasini, Kalyani Peri, Gary Cutter, Darcy A. Krueger, and the PREVeNT Study Group. Early treatment with vigabatrin does not decrease focal seizures or improve cognition in tuberous sclerosis complex: The PREVeNT trial. Annals of Neurology, 2023. ISSN 0364-5134. doi: 10.1002/ana.26778.

95 Tommaso Fedele, Maryse van ‘t Klooster, Sergey Burnos, Willemiek Zweiphenning, Nicole van Klink, Frans Leijten, Maeike Zijlmans, and Johannes Sarnthein. Automatic detection of high frequency oscillations during epilepsy surgery predicts seizure outcome. Clinical Neurophysiology, 127(9):3066–3074, 2016. ISSN 1388-2457. doi: 10.1016/j.clinph.2016.06.009.

96 Maryse A. van ‘t Klooster, Nicole E.C. van Klink, Willemiek J.E.M. Zweiphenning, Frans S.S. Leijten, Rina Zelmann, Cyrille H. Ferrier, Peter C. van Rijen, Willem M. Otte, Kees P.J. Braun, Geertjan J.M. Huiskamp, and Maeike Zijlmans. Tailoring epilepsy surgery with fast ripples in the intraoperative electrocorticogram. Annals of Neurology, 81(5):664–676, 2017. ISSN 0364-5134. doi: 10.1002/ana.24928.

97 Ece Boran, Johannes Sarnthein, Niklaus Krayenbühl, Georgia Ramantani, and Tommaso Fedele. High-frequency oscillations in scalp EEG mirror seizure frequency in pediatric focal epilepsy. Scientific Reports, 9(1):16560, 2019. doi: 10.1038/s41598-019-52700-w.

98 Eline V. Schaft, Dongqing Sun, Maryse A. van ‘t Klooster, Dorien van Blooijs, Paul L. Smits, Willemiek J.E.M. Zweiphenning, Peter H. Gosselaar, Cyrille H. Ferrier, Maeike Zijlmans, and on behalf of the RESPect database study group. Spatial and temporal properties of intra-operatively recorded spikes and high frequency oscillations in focal cortical dysplasia. Clinical Neurophysiology, 162:210–218, 2024. ISSN 1388-2457. doi: 10.1016/j.clinph.2024.03.038.

99 Andrea Giovannucci, Johannes Friedrich, Pat Gunn, Jérémie Kalfon, Brandon L Brown, Sue Ann Koay, Jiannis Taxidis, Farzaneh Najafi, Jeffrey L Gauthier, Pengcheng Zhou, Baljit S Khakh, David W Tank, Dmitri B Chklovskii, and Eftychios A Pnevmatikakis. CaImAn an open source tool for scalable calcium imaging data analysis. eLife, 8:e38173, 2019. doi: 10.7554/elife.38173.

